# Intracellular targeting of a light-activated GPCR elicits spatially selective signalling

**DOI:** 10.64898/2026.01.29.702683

**Authors:** Chantel Mastos, Christina G Gangemi, Alexandra-Madelaine Tichy, Bonan Liu, Cameron Nowell, Andrew M Ellisdon, Harald Janovjak, Michelle L Halls

**Author notes:** Correspondence to: Associate Professor Michelle L Halls. Department of Genetics, Cell Biology and Development, University of Minnesota, Minneapolis, USA.

## Abstract

G protein-coupled receptors (GPCRs) initiate unique cell responses dependent on their sub-cellular location. Major advances in our understanding of sub-cellular GPCR signalling have been driven by experimental approaches that manipulate receptor trafficking, block signalling at specific intracellular sites, or compare receptor signalling in response to ligands with differing membrane permeability. Here, we describe an optogenetic platform that allows a single stimulus to activate the same GPCR at different intracellular membranes, enabling direct comparisons of spatial signalling in intact cells. To achieve this, we have directed a light-activated GPCR chimera to endosomal, mitochondrial, Golgi and nuclear membranes using C-terminal targeting sequences. We demonstrate activation of cAMP and ERK signalling in response to light at all sites; however, each targeted receptor generates a unique transcriptional fingerprint dependent on its sub-cellular location. Our engineered optogenetic platform reveals that GPCRs targeted to distinct intracellular locations are self-sufficient signalling platforms and highlights GPCR positioning as a critical driver of signalling.

**Teaser:** Controlling GPCR signals by light and location reveals how intracellular positioning shapes cellular responses.

## Introduction

G protein-coupled receptors (GPCRs) are the largest family of transmembrane proteins in the human genome. They transduce inputs from a diverse range of stimuli (including light, odour, hormones, and peptides) to allow the cell to dynamically respond to its environment (*1*). The restriction of GPCR signalling to discrete microdomains – or spatial compartmentalisation of signalling – is a mechanism that allows the >800 GPCRs to couple to a finite set of intracellular signalling pathways but initiate distinct cellular responses (*2*). The idea that GPCR signalling is compartmentalised initially arose from studies of the activation of β-adrenoceptors and prostaglandin receptors in cardiomyocytes; both receptors increase cAMP in these cells, but only the cAMP generated by the β-adrenoceptor increases myocyte contractility (*3-6*).

The cellular consequences of GPCR signalling change depending on receptor location. Some receptors induce distinct responses at the plasma membrane compared to after their internalisation and trafficking to intracellular organelles such as the endosomal network (*7-11*) and the Golgi apparatus (*12-15*). For example, sustained signalling of the neurokinin 1 receptor from endosomes (but not at the plasma membrane) is necessary for pain transmission (*8, 11*). In addition, the β_2_-adrenoceptor (β_2_AR) regulates distinct genes depending on whether the receptor is at the plasma membrane or internalised to endosomes (*10*). In particular, the positioning of endosomes containing the β_2_AR relative to the nucleus is important (*16*), as gene transcription is controlled by the propagation of endosomal β_2_AR signalling to the nuclear compartment (*16, 17*). These studies (and many others) have provided a fundamental understanding of how GPCR location controls cellular responses. They have principally depended on global blockade of endocytosis (which is critical for a broad array of cellular functions) or comparisons of signalling in response to cell permeable versus impermeable ligands (thus introducing the potential for ligand bias and differences in ligand efficacy). Recent studies have addressed these challenges using cellular engineering to artificially relocate endosomes (*16*) or to selectively block GPCRs at discrete intracellular sites using targeted nanobodies (*17*). When considered all together, these orthogonal strategies provide complementary but distinct insights into how receptor trafficking from the plasma membrane to different intracellular sites changes GPCR signalling.

In parallel to studies investigating intracellular GPCR signalling initiated at the plasma membrane, other work has focused on GPCRs that natively reside within intracellular compartments. Some GPCRs are found at the Golgi, nucleus or mitochondria, where they initiate signalling *in situ* and trigger cellular responses distinct from those activated when the same receptor is stimulated at the plasma membrane (*13-15, 18-21*). For example, in cardiomyocytes the β_1_-adrenoceptor (β_1_AR) is distributed both at the cell surface and at the Golgi, and receptor activation leads to spatially distinct cAMP pools (*13, 22*); cell surface β_1_AR increases contractile force while Golgi β_1_AR increases the rate of relaxation and cell hypertrophy (*14, 15*). Inspired by these observations of intracellular GPCR signalling *in situ*, we asked what if we pre-emptively manipulate the cellular location of a GPCR prior to stimulation? This would allow us to compare responses across cells where the only difference is the starting location of the receptor. There are two principal challenges in developing such a model system. First, we need to ensure that receptors at the plasma membrane and at each intracellular location have equivalent access to the activating stimulus. Second, we need to identify appropriate organelle targeting sequences: while there are many well-defined targeting motifs for soluble, cytosolic proteins, such sequences are not established for transmembrane proteins.

The use of light *in lieu* of a ligand overcomes many obstacles associated with ligand transport to intracellular locations – the delivery of light is unaffected by diffusion or membrane permeability, and an exogenously introduced light-sensitive receptor can be specifically activated without any coincident stimulation of endogenous receptor populations. We therefore developed a model system based on the reengineered opto-β_2_-adrenoceptor (opto-β_2_AR^2.0^), a second-generation optogenetic GPCR (optoGPCR) chimera modified for increased G protein binding (*23*). The opto-β_2_AR^2.0^ combines the extracellular, transmembrane domains and intracellular loop (ICL) 1 of the light-activated rhodopsin with the intracellular (ICL2, ICL3, C-terminus) and G protein-interacting regions of transmembrane domains 3-6 of the hamster β_2_AR (*24-28*) (Figure 1A). The β_2_AR is a very well-characterised GPCR that is activated by adrenaline and noradrenaline. This receptor can signal *via* cAMP at the plasma membrane and *via* cAMP and ERK pathways following internalisation to endosomes (*9, 17*). The opto-β_2_AR retains fidelity of G protein signalling compared to the β_2_AR; it can activate both cAMP and ERK phosphorylation signalling pathways, and internalise following activation (*24, 25, 27*).

**Figure 1.**
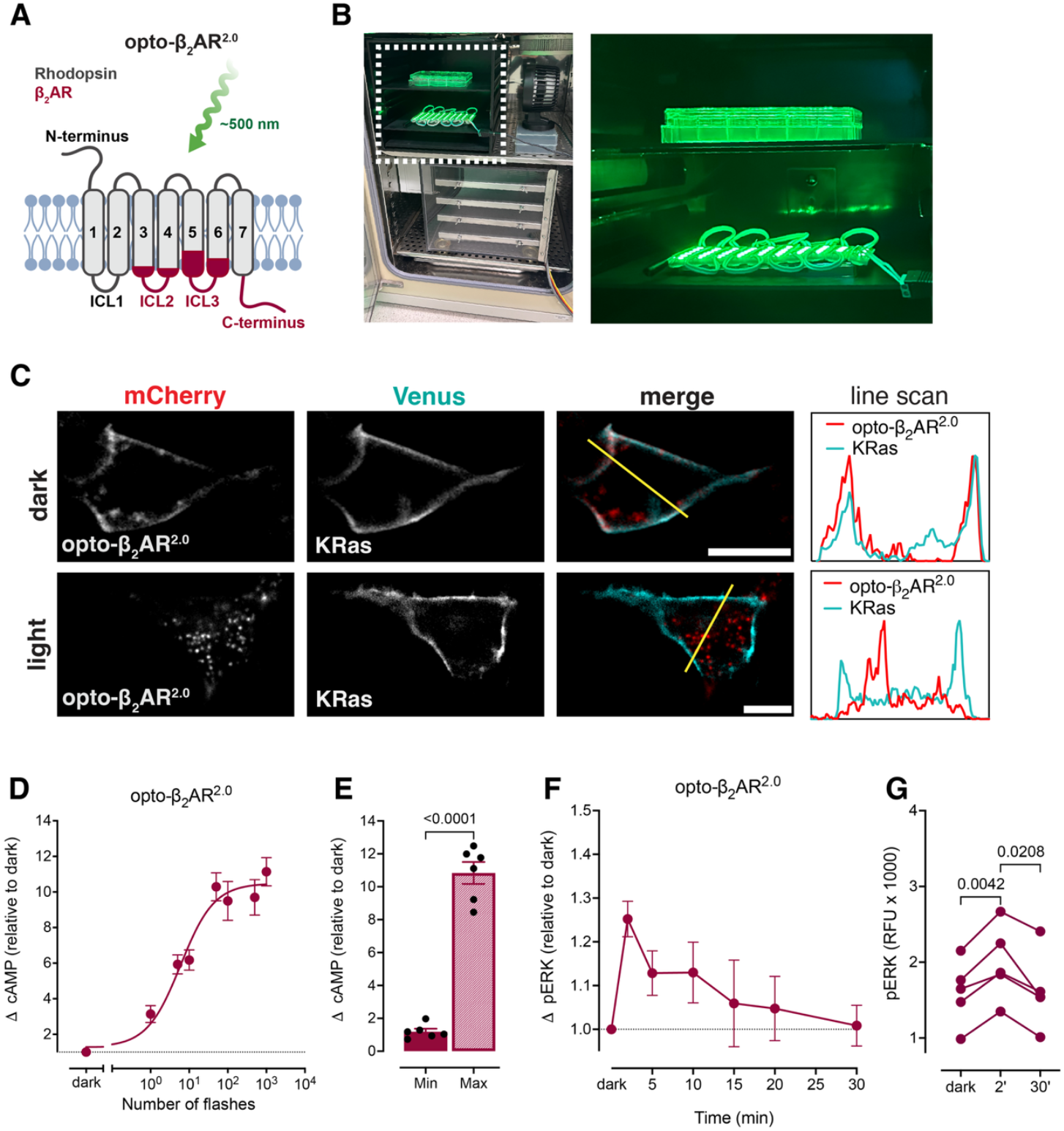
Light activation of the opto-β_2_AR^2.0^ internalises the receptor, and increases cAMP and ERK signalling. **A)** The chimeric opto-β_2_AR^2.0^ contains the light-activated transmembrane (TM, domains are numbered), ICL1 and extracellular domains of rhodopsin (shown in grey), and the G protein-binding regions of TM3-6 and intracellular regions (ICL2, ICL3, C-terminal tail) of the hamster β_2_AR (shown in maroon). The opto-β_2_AR^2.0^ can be activated by green LEDs using a **B**) modular LED shelving system that stacks in a standard cell culture incubator (upper shelf; lower shelf holds a hypoxia chamber). Inset shows close view of the LED array and 6-well plate in the shelving system. **C**) HEK293 cells were co-transfected with the plasma membrane marker KRas-Venus and the mCherry-opto-β_2_AR^2.0^, then either fixed in the dark or live cells were activated by light. Representative images show the opto-β_2_AR^2.0^ (left panel), KRas-Venus plasma membrane marker (middle panel), and the merged image (right panel), with the yellow line indicating the region analysed in the line scan intensity graph (far right panel). Scale bars are 10 μm. **D**) Light-induced change in cAMP expressed relative to dark in HEK293 cells transfected with the opto-β_2_AR^2.0^. Symbols represent means, and error bars show SEM (n=6). **E**) Minimum and maximum cAMP responses calculated from the light-response curve shown in D. Symbols show values from independent experiments, bars show mean, and error bars SEM (n=6); p-value determined by paired t-test. **F**) Light-induced increase in ERK phosphorylation relative to dark in HEK293 cells transfected with the opto-β_2_AR^2.0^. Symbols represent means and error bars show SEM (n=5). **G**) Light caused a transient increase in ERK phosphorylation with a peak at 2 min, that decreased by 30 min. Symbols show relative fluorescence units (RFU) from independent experiments, p-value determined by repeated measures one-way ANOVA with Dunnett’s multiple comparisons test.

Here, we first established our sub-cellular targeted opto-β_2_AR^2.0^ toolbox using C-terminal intracellular targeting sequences. We then used our synthetic system to demonstrate that activation of the same receptor by the same stimulus from distinct cellular locations induces a unique transcriptional fingerprint.

## Results

### Activation of the opto-β_2_AR^2.0^ mimics wild-type β_2_AR signalling

We first confirmed that the opto-β_2_AR^2.0^ fused to an N-terminal mCherry tag retains G protein signalling of the wild-type β_2_AR in our HEK293 cell system. The opto-β_2_AR^2.0^ is activated in response to green light (Figure 1A) that can be delivered using microscopes, plate readers, or a custom-built modular light-emitting diode (LED) shelving system (*29*) (Figure 1B). Confocal imaging of cells transfected with the opto-β_2_AR^2.0^ that were fixed in the dark showed that the receptor was localised to the plasma membrane, as determined using a plasma membrane marker, KRas-Venus (Figure 1C). There was also a small proportion of mCherry fluorescence observed in intracellular punctae. This is consistent with constitutive receptor recycling, which is also observed for the wild-type β_2_AR (*30*). Imaging of live cells (in which light from the microscope activated the receptor) showed internalisation of the opto-β_2_AR^2.0^ away from the plasma membrane into intracellular punctae (Figure 1C). This also mimics the activation of the wild-type β_2_AR by ligand, whereby the receptor undergoes β-arrestin-dependent internalisation to endosomes (*31, 32*).

Next, we confirmed that green light stimulation of the opto-β_2_AR^2.0^ could activate the expected cAMP and ERK intracellular signalling pathways. Application of an increasing number of light flashes showed a light-dependent increase in cAMP accumulation (Figure 1D-E). Stimulation of the opto-β_2_AR^2.0^ also caused a transient increase in ERK phosphorylation, which peaked at 2 min and returned to baseline levels by ∼20 min (Figure 1F-G).

### C-terminal sequences can target the opto-β_2_AR^2.0^ to organelles

To target the opto-β_2_AR^2.0^ to specific intracellular membranes, we performed a literature search for C-terminal sequence motifs important for retaining a variety of transmembrane proteins at organelles. We identified at least two sequences for the targeting of transmembrane proteins to early endosomes, the endoplasmic reticulum, the Golgi, mitochondria, and the nucleus (Table 1; see Supplementary Table 1 for a complete list of identified sequences). Each targeting sequence was separately added to the C-terminus of the opto-β_2_AR^2.0^. A plasma membrane-restricted variant of the first generation opto-β_2_AR was generated by mutating C-terminal serine residues to prevent GRK-mediated receptor phosphorylation, β-arrestin binding and receptor internalisation (*25*). However, we reasoned that the removal of these intracellular phosphorylation sites may also restrict or alter receptor-effector interactions and inadvertently influence the signalling profile. We therefore used the untargeted opto-β_2_AR^2.0^ for comparisons.

**Table 1.**
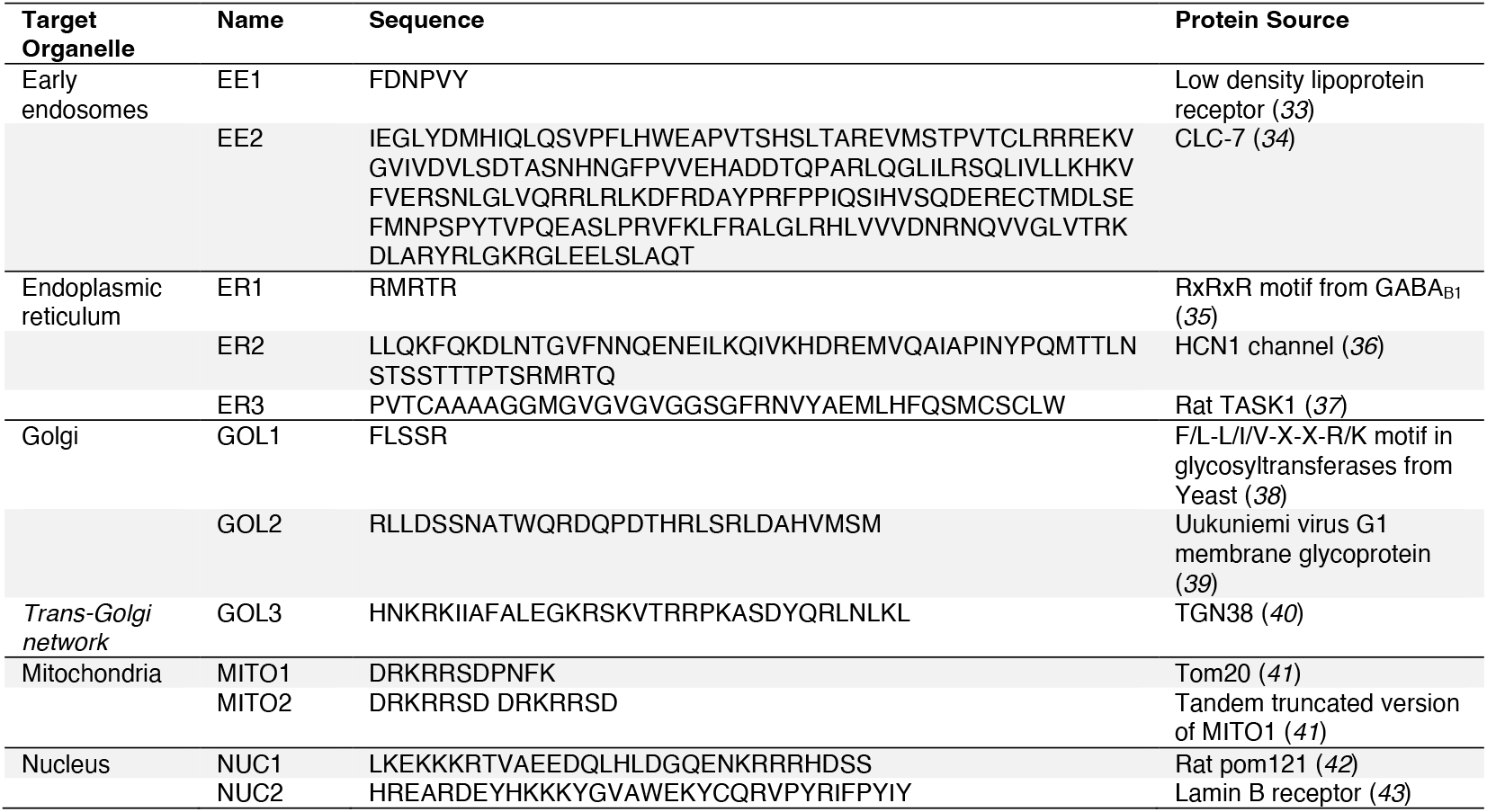
C-terminal organelle targeting sequences. Sequences that were identified and tested for organelle targeting of transmembrane proteins.

To determine if the identified sequences consistently localised the transiently transfected opto-β_2_AR^2.0^ to the target organelle, we developed a semi-automated analysis pipeline. Each sub-cellular region or organelle was defined by the transfection of a well-characterised Venus-tagged marker (KRas-Venus for plasma membrane, Rab5a-Venus for early endosomes, Venus-PTP1b for the endoplasmic reticulum, Venus-giantin for the Golgi, Venus-mito-7 for mitochondria), or stain (Hoescht DNA stain for the nucleus). The amount of mCherry fluorescence from the targeted opto-β_2_AR^2.0^ contained within the same pixels as the organelle marker was quantified. Receptors with C-terminal targeting sequences were short-listed as correctly localised if we observed co-incident fluorescence with the organelle marker in >80% of cells that were co-transfected with the two proteins, with minimal variation in this localisation over the cell population (Supplementary Figure 1A-D). We then confirmed this localisation with at least one additional marker using the same criteria but a different expression or detection system (e.g. transient transfection of a fluorescently-tagged marker followed by immunostaining of an endogenous marker protein).

To target transmembrane proteins to the early endosome, we tested two C-terminal sequences, which we named EE1 and EE2 (Table 1). Sequence EE2, but not EE1, consistently localised the opto-β_2_AR^2.0^ to early endosomes as defined by the Rab5a marker, with 84.70±1.67% (mean±SEM from three independent experiments; 45 cells total) of the mCherry fluorescence contained in the same pixels as the Venus marker (Figure 2A-B, Supplementary Figure 2A-E). To confirm the localisation of the EE2 fusion opto-β_2_AR^2.0^ in endosomes, we performed immunostaining using an anti-EEA1 antibody (Figure 2A-B, Supplementary Figure 2E-G). Despite the mCherry signal appearing as intracellular punctae, within the cell population we observed a large variation in the amount of mCherry fluorescence which coincided with EEA1 immunofluorescence. To investigate this, we classified the opto-β_2_AR^2.0^ as residing in EEA1-negative or EEA1-positive punctae (using 40% or 60% co-location as a cut-off, respectively; this removed 12 cells from the analysis). We found that the opto-β_2_AR^2.0^ was distributed across EAA1-positive and EEA1-negative endosomes (EEA1-positive: 86.61±3.37% across 55 cells; EEA1-negative: 19.80±4.39% across 51 cells; mean±SEM from three independent experiments; Supplementary Figure 2F-G). Our observations are consistent with a previous systematic study of structures labelled by Rab5a and EEA1; Rab5a predominantly marks small endocytic vesicles and early endosomes, whereas EEA1 identifies a sub-population of Rab5a-positive early endosomes but also labels Rab5a-negative late endosomes (*44*). As such, the EE2 fusion appears to direct the opto-β_2_AR^2.0^ to Rab5a-positive endosomes, with no preference for the EEA1-positive sub-population. We therefore termed the EE2 fusion opto-β_2_AR^2.0^-endo (early endosome localised opto-β_2_AR^2.0^).

**Figure 2.**
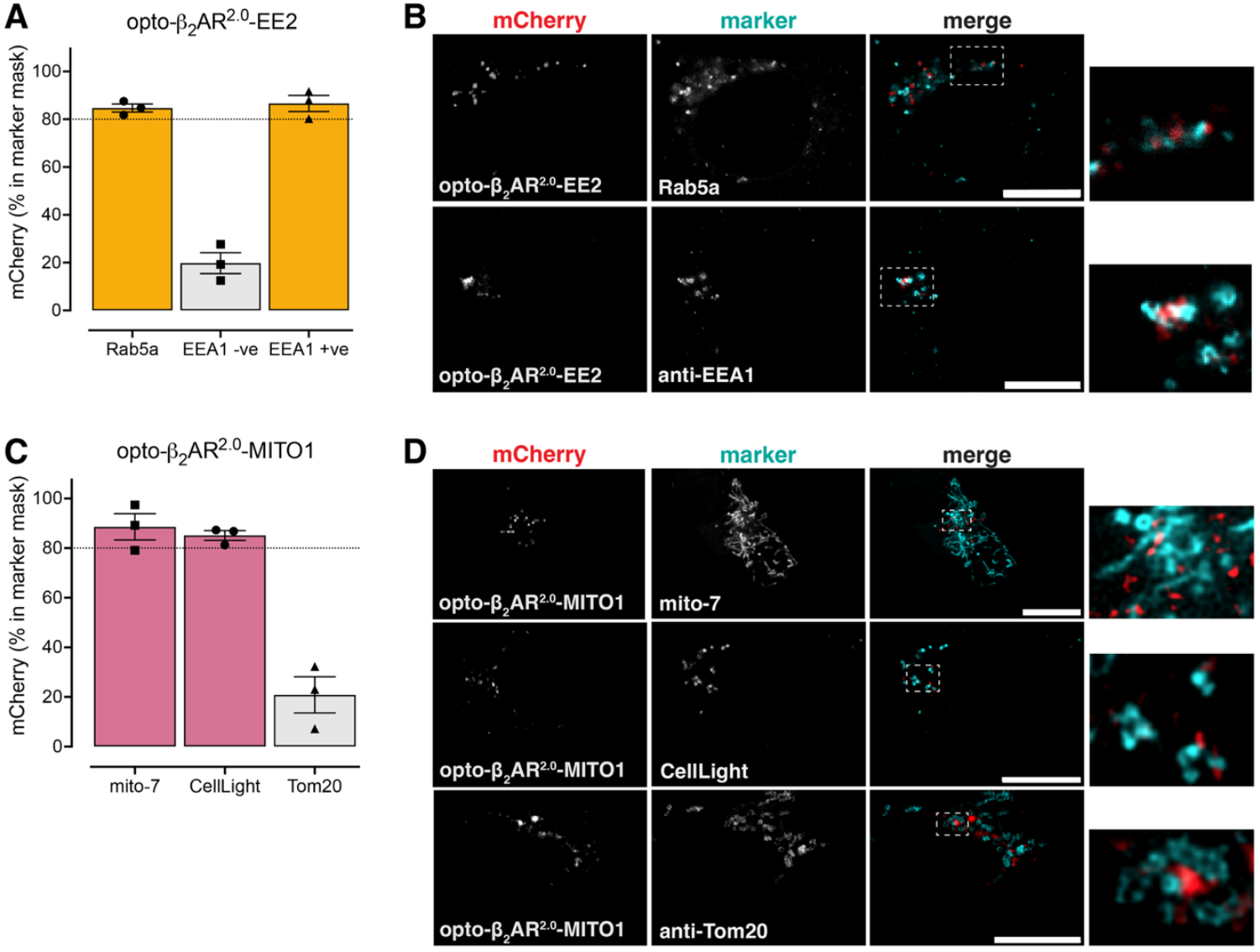
C-terminal sequences can target the opto-β_2_AR^2.0^ to early endosomes and mitochondria. C-terminal targeting sequences were added to the opto-β_2_AR^2.0^ to localise the receptor to **A-B**) early endosomes, or **C-D**) mitochondria. HEK293 cells were transfected with the targeted mCherry-opto-β_2_AR^2.0^ and either co-transfected with a Venus-tagged intracellular marker, infected with a GFP-tagged CellLight marker, or immunostained with an antibody against an organelle resident protein. The amount of mCherry fluorescence within the pixel area defined by intracellular markers was quantified (n=3). For bar graphs (A and C), symbols show median values from all cells from each biological replicate (shown in Supplementary Figure 2D, 2F-G, 3C, 3F-G), the bar shows the mean and error bars show SEM (n=3). “EEA1 -ve” and “EEA1 +ve” refer to mCherry punctae that were negative or positive for EEA1 immunostaining, respectively. For image panels (B and D), representative images show mCherry-opto-β_2_AR^2.0^ (left panel), Venus- or GFP-tagged marker or immunostaining (middle panel), and merged image (right panel). The inset (far right) shows the region boxed in white. Scale bars are 10 μm.

We next tested two C-terminal sequences to target the opto-β_2_AR^2.0^ to the mitochondria, MITO1, and a shortened version of the same sequence as a tandem repeat, MITO2 (Table 1). Sequence MITO1, but not MITO2, consistently localised the opto-β_2_AR^2.0^ to the mitochondria as defined by the mito-7 marker, with 88.57±5.30% (mean±SEM from 3 independent experiments; 32 cells total) of the mCherry fluorescence contained within the same pixels as the Venus marker (Figure 2C-D and Supplementary Figure 3A-E). To confirm the localisation of the opto-β_2_AR^2.0^ MITO1 fusion to the mitochondria, we used an additional two approaches: CellLight Mito-GFP (baculovirus containing GFP-labelled leader sequence of EF1α pyruvate dehydrogenase) and immunostaining using an anti-Tom20 antibody. We found that the opto-β_2_AR^2.0^ resided within the region defined by CellLight Mito-GFP fluorescence (85.08±1.92%, mean±SEM from 3 independent experiments; 35 cells total), but not the region defined by Tom20 immunofluorescence (20.85±7.3 2%, mean±SEM from 3 independent experiments; 70 cells total) (Figure 2C-D and Supplementary Figure 3E-G). The three protein markers reside on different mitochondrial membranes. The mito-7 marker uses the leader sequence of COX8A which faces the mitochondrial matrix on the inner mitochondrial membrane (*45*). Similarly, CellLight Mito-GFP localises to the inner mitochondrial membrane with the GFP-tag in the mitochondrial matrix (*46*). In contrast, the anti-Tom20 antibody binds to the cytosolic face of Tom20 which resides in the outer mitochondrial membrane (*47*). Together, this suggests that the MITO1 sequence directs the opto-β_2_AR^2.0^ to the inner mitochondrial membrane. We therefore termed the MITO1 fusion opto-β_2_AR^2.0^-mito (mitochondria localised opto-β_2_AR^2.0^).

**Figure 3.**
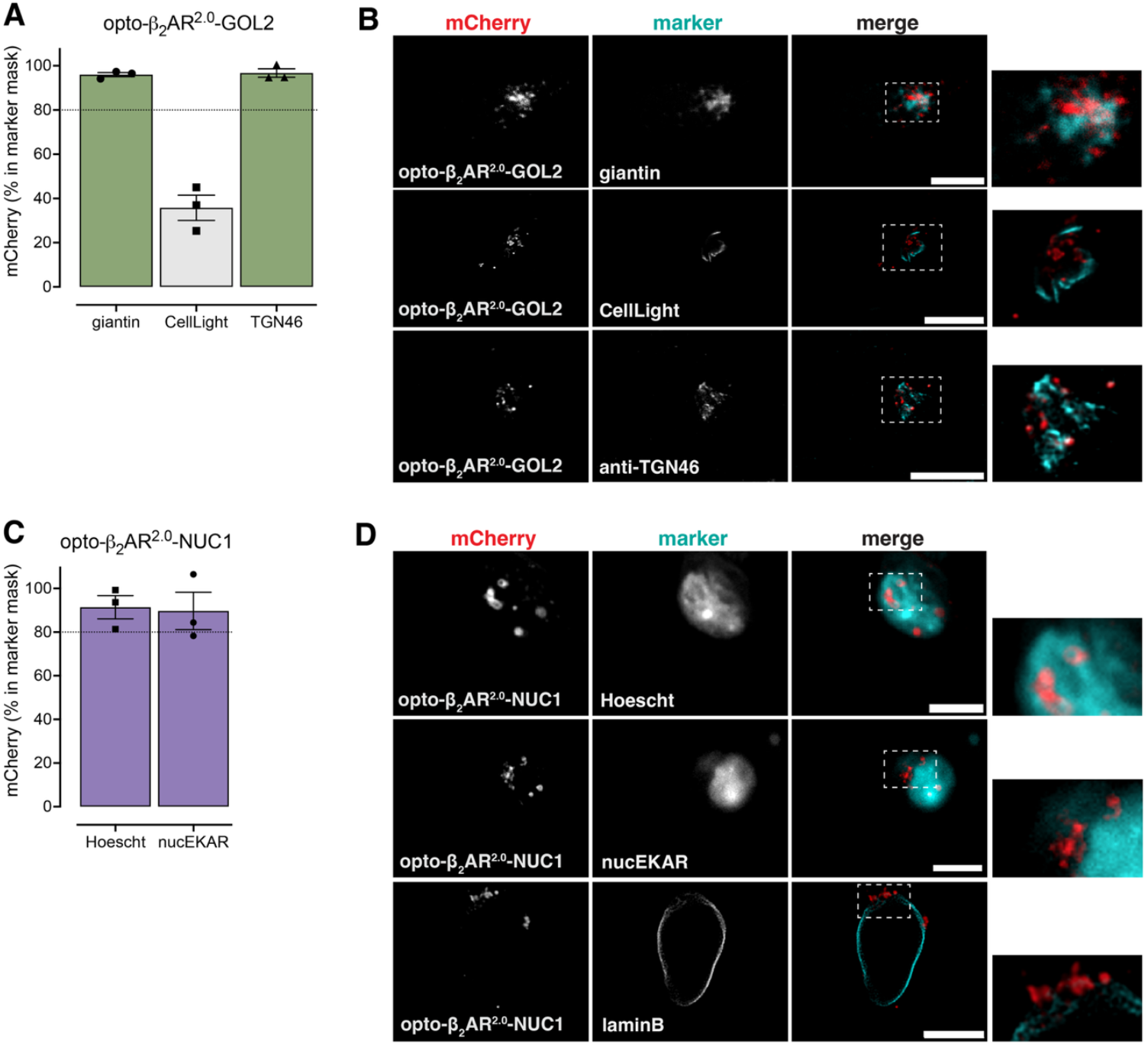
C-terminal sequences can target the opto-β_2_AR^2.0^ to the Golgi, and the nucleus. C-terminal targeting sequences were added to the opto-β_2_AR^2.0^ to localise the receptor to the **A-B**) Golgi, or **C-D**) nucleus. HEK293 cells were co-transfected with the targeted mCherry-opto-β_2_AR^2.0^ and either co-transfected with a Venus-tagged intracellular marker, infected with a GFP-tagged CellLight marker, immunostained with an antibody against an organelle resident protein, or incubated with the Hoescht DNA stain. The amount of mCherry fluorescence within the pixel area defined by intracellular markers was quantified (n=3). For bar graphs (A and C), symbols show median values from all cells from each biological replicate (shown in Supplementary Figures 4C-E, 4G-H, 5C-D, 5F), the bar shows the mean and error bars show SEM (n=3). For image panels (B and D), representative images show mCherry-opto-β_2_AR^2.0^ (left panel), Venus- or GFP-tagged marker or immunostaining or Hoescht stain (middle panel), and merged image (right panel). Inset (far right) shows area boxed in white. Scale bars are 10 μm.

To target the opto-β_2_AR^2.0^ to the Golgi we tested three C-terminal sequences, named GOL1, GOL2 and GOL3 (Table 1). Only sequence GOL2 consistently localised the opto-β_2_AR^2.0^ to the Golgi as defined by the giantin marker, with 95.99±0.93% (mean±SEM from three independent experiments; 39 cells total) of the mCherry fluorescence contained within the same pixels as the Venus marker (Figure 3A-B, Supplementary Figure 4A-F). To confirm the localisation of the opto-β_2_AR^2.0^ GOL2 fusion to the Golgi, we used two additional approaches: CellLight Golgi-GFP (baculovirus containing GFP-labelled N-acetylgalactosaminyltransferase 2) and immunostaining using an anti-trans-Golgi network protein 2 (TGN46) antibody. While we found that the opto-β_2_AR^2.0^ consistently resided within the region defined by TGN46 immunofluorescence (96.70±1.93%, mean±SEM from three independent experiments; 51 cells total), there was minimal overlap with the CellLight Golgi-GFP fluorescence (35.78±5.71%, mean±SEM from three independent experiments, 35 cells total) (Figure 3A-B and Supplementary Figure 4F-H). The three protein markers we used have differential localisation within the Golgi; Venus-giantin (residues 3131-3259 of giantin) is principally found within the peripheral membranes of the medial-Golgi (*48*) and the trans-Golgi network (*49*), N-acetylgalactosaminyltransferase 2 is located to the medial/trans-Golgi within the interior (*48*), and TGN46 defines the trans-Golgi network (*50*). Taken together, this suggests GOL2 is targeting the opto-β_2_AR^2.0^ to peripheral trans-Golgi and the trans-Golgi network. We therefore termed the GOL2 fusion opto-β_2_AR^2.0^-golgi (Golgi localised opto-β_2_AR^2.0^).

Two C-terminal sequences were tested for nuclear targeting of the opto-β_2_AR^2.0^, NUC1 and NUC2 (Table 1). Sequence NUC1, but not NUC2, consistently localised the opto-β_2_AR^2.0^ to the nucleus as defined by the Hoescht DNA stain, with 91.40±5.30% (mean±SEM from three independent experiments; 47 cells total) of the mCherry fluorescence contained within the same pixels as the Hoescht stain (Figure 3C-D and Supplementary Figure 5A-E). Localisation of the opto-β_2_AR^2.0^ NUC1 fusion to the nucleus was confirmed by co-transfecting cells with a nuclear localised ERK biosensor, nucEKAR (*51*) (89.70 ±5.30% of mCherry fluorescence within the nucEKAR region, mean±SEM from three independent experiments, 47 cells total) (Figure 3C-D and Supplementary Figure 5E-F). As neither Hoescht nor nucEKAR are membrane-associated, we further confirmed the localisation of the targeted opto-β_2_AR^2.0^ to the nuclear membrane using a Venus-tagged resident protein of the inner nuclear membrane, lamin B (Figure 3C-D and Supplementary Figure 5E). We termed the NUC1 fusion opto-β_2_AR^2.0^-nuc (nuclear localised opto-β_2_AR^2.0^).

Finally, three C-terminal sequences were tested for targeting of the opto-β_2_AR^2.0^ to the endoplasmic reticulum, named ER1, ER2 and ER3 (Table 1). None of these sequences allowed the selective retention of the opto-β_2_AR^2.0^ in the endoplasmic reticulum, with a small degree of plasma membrane localisation always evident (Supplementary Figure 6A-F). We therefore abandoned efforts to target the opto-β_2_AR^2.0^ to the endoplasmic reticulum, and proceeded with the opto-β_2_AR^2.0^ targeted to the early endosomes, mitochondria, Golgi, and nucleus.

### Light activation of targeted opto-β_2_AR^2.0^s increases cAMP and pERK

We then asked whether each targeted opto-β_2_AR^2.0^ can signal *in situ* via cAMP and ERK pathways, without a requirement to traffic to this location from the plasma membrane (Figures 4 and 5). Light-mediated cAMP accumulation or ERK phosphorylation was quantified using cell population immunoassays that measured the global signal throughout the cell. For all targeted receptors (early endosomes, mitochondria, Golgi, and nucleus), increasing amounts of light caused an increase in cAMP accumulation (Figure 4A-H). Similarly, light stimulation caused an increase in ERK phosphorylation for all intracellular receptors (Figure 5A-H).

**Figure 4.**
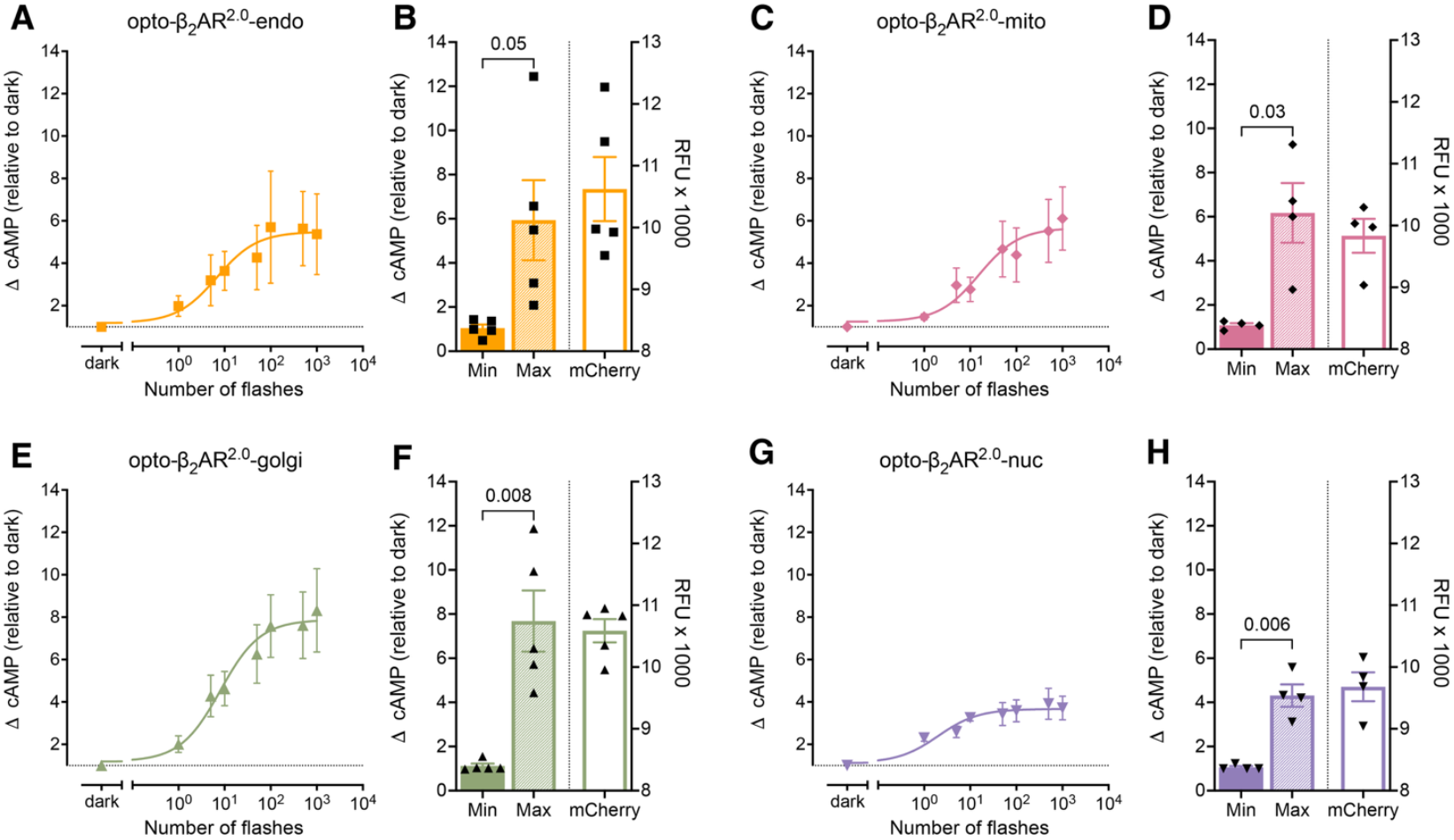
Light activation of the targeted opto-β_2_AR^2.0^s increases cAMP accumulation from early endosomes, mitochondria, Golgi, and the nucleus. cAMP was measured in HEK293 cells transiently transfected with **A-B**) opto-β_2_AR^2.0^-endo (n=5), **C-D**) opto-β_2_AR^2.0^-mito (n=4), **E-F**) opto-β_2_AR^2.0^-golgi (n=5), and **G-H**) opto-β_2_AR^2.0^-nuc (n=4) following exposure to increasing amounts of light. For light-response curves (A, C, E, G) symbols represent means and error bars show SEM. Bar graphs (B, D, F, H) display the minimum and maximum values calculated from the fit of the light-response curves for each individual experiment (left y-axis); receptor expression levels were simultaneously estimated by quantifying the mCherry fluorescence (relative fluorescence units, RFU) in each well (right y-axis). Symbols show values from independent experiments, bars show mean and error bars show SEM, p-values calculated by paired t-test.

**Figure 5.**
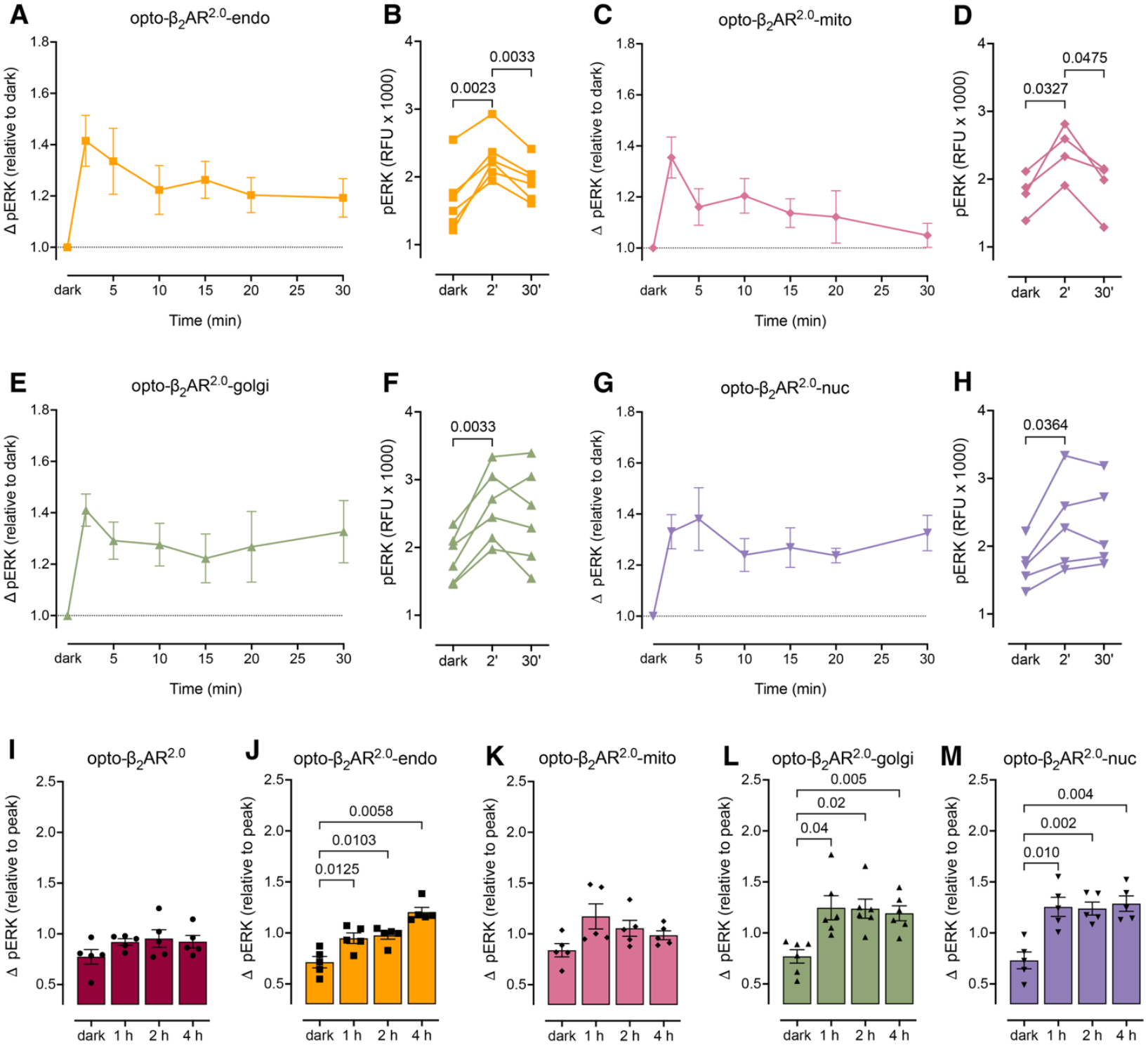
Light activation of the targeted opto-β_2_AR^2.0^s increases ERK phosphorylation from early endosomes, mitochondria, Golgi, and the nucleus. ERK phosphorylation was measured in HEK293 cells transiently transfected with **A-B**) opto-β_2_AR^2.0^-endo (n=6), **C-D**) opto-β_2_AR^2.0^-mito (n=4), **E-F**) opto-β_2_AR^2.0^-golgi (n=6), and **G-H**) opto-β_2_AR^2.0^-nuc (n=5) following exposure to light over a 30 min time course. For time courses (A, C, E, G) symbols represent means and error bars show SEM. Bar graphs (B, D, F, H) display the ERK phosphorylation in relative fluorescence units (RFU) in the dark, and after 2 min or 30 min exposure to light. Symbols show values from independent experiments, p values calculated using repeated measures one-way ANOVA with Dunnett’s multiple comparisons test. To confirm whether increases in ERK phosphorylation were transient or sustained, the impact of prolonged light stimulation (up to 4 h) of **I**) opto-β_2_AR^2.0^ (n=5), **J**) opto-β_2_AR^2.0^-endo (n=5), **K**) opto-β_2_AR^2.0^-mito (n=5), **L)** opto-β_2_AR^2.0^-golgi (n=6), or **M**) opto-β_2_AR^2.0^-nuc (n=5) was measured. Symbols show values from independent experiments, bars show mean and error bars show SEM, p values calculated using one-way ANOVA with Dunnett’s multiple comparisons test.

Although the targeted receptors were all capable of activating cAMP and ERK signalling in response to light, the magnitude and the temporal dynamics of these responses varied based on location. Activation of the opto-β_2_AR^2.0^-endo, opto-β_2_AR^2.0^-golgi and opto-β_2_AR^2.0^-mito by light resulted in maximal cAMP responses that were not statistically different from that of the untargeted opto-β_2_AR^2.0^ (Figure 1D-E, Figure 4A-F and Supplementary Figure 7A). In contrast, activation of opto-β_2_AR^2.0^-nuc produced a smaller amount of cAMP accumulation compared to the untargeted opto-β_2_AR^2.0^ (Figure 1G-H, Supplementary Figure 7A). This difference in the magnitude of the cAMP response was not due to differences in receptor expression levels in the cell, as approximated by the intensity of mCherry fluorescence (Figure 4B, D, F, H, Supplementary Figure 7B-C). As such, receptor location appears to dictate the magnitude of the cAMP response, with smaller amounts of cAMP generated by the receptor localised to the nucleus.

Despite no influence of receptor location on the magnitude of the initial peak ERK phosphorylation response at 2 min (Supplementary Figure 7D), we found that the intracellular location dictated different temporal profiles of ERK phosphorylation (Figure 5A-H and Supplementary Figure 7E-F). Light activation of opto-β_2_AR^2.0^-endo induced a transient increase in ERK phosphorylation which peaked at 2 min and then declined to a sustained, steady-state level above baseline (Figure 5A-B, Supplementary Figure 7E). We calculated the fold change in the response at 30 min relative to the response at 2 min to yield a response ratio. A response ratio of 1 indicates an equal magnitude of ERK phosphorylation at the two time points, and therefore a sustained response, while a response ratio less than 1 indicates a smaller response at 30 min compared to 2 min, representing a transient response. As expected, the response ratios for opto-β_2_AR^2.0^-endo and the untargeted opto-β_2_AR^2.0^ were significantly below 1 (Supplementary Figure 7F). When we activated opto-β_2_AR^2.0^-mito with light we observed a similar transient increase in ERK phosphorylation that declined back to baseline by 30 min, with a response ratio below 1 (Figure 5C-D and Supplementary Figure 7E-F). In contrast, activation of both the opto-β_2_AR^2.0^-golgi and the opto-β_2_AR^2.0^-nuc caused a sustained increase in ERK phosphorylation which remained elevated over the 30 min time course, with response ratios that were not different from 1 (Figure 5E-H and Supplementary Figure 7E-F).

We then conducted a longer light stimulation time course over 4 hours (Figure 5I-M). These experiments confirmed that the untargeted opto-β_2_AR^2.0^ (Figure 5I) and opto-β_2_AR^2.0^-mito (Figure 5K) induced only transient increases in ERK phosphorylation, with no effect of light activation for 1 hour, 2 hours, or 4 hours compared to dark control. In contrast, light activation of opto-β_2_AR^2.0^-endo (Figure 5J), opto-β_2_AR^2.0^-golgi (Figure 5L) and opto-β_2_AR^2.0^-nuc (Figure 5M) resulted in a sustained increase in ERK phosphorylation that remained elevated for up to 4 hours (Supplementary Figure 7G). Similar to cAMP signalling, these distinct temporal profiles of ERK phosphorylation were not due to any differences in receptor expression (Supplementary Figure 7H-I). Together, these data demonstrate that the intracellular location of the opto-β_2_AR^2.0^ regulates both the magnitude of signalling and the timescale on which this occurs.

### Targeted opto-β_2_AR^2.0^s induce distinct gene expression profiles

The wild-type β_2_AR regulates distinct genes depending on whether the receptor is at the plasma membrane or internalised to endosomes, and on the positioning of these endosomes relative to the nucleus (*10, 16*). We therefore used our engineered system to determine whether *in situ* activation of the opto-β_2_AR^2.0^ can impact gene transcription from organelles other than the endosomes. Light-induced changes in gene expression following activation of the opto-β_2_AR^2.0^ at each intracellular location were quantified by RNA-seq in stably transfected HEK293 cells (Figure 6, Supplementary Figure 8A-E). Across four biological replicates, 27,775 known genes were successfully mapped to gene IDs (Supplementary Table 2).

**Figure 6.**
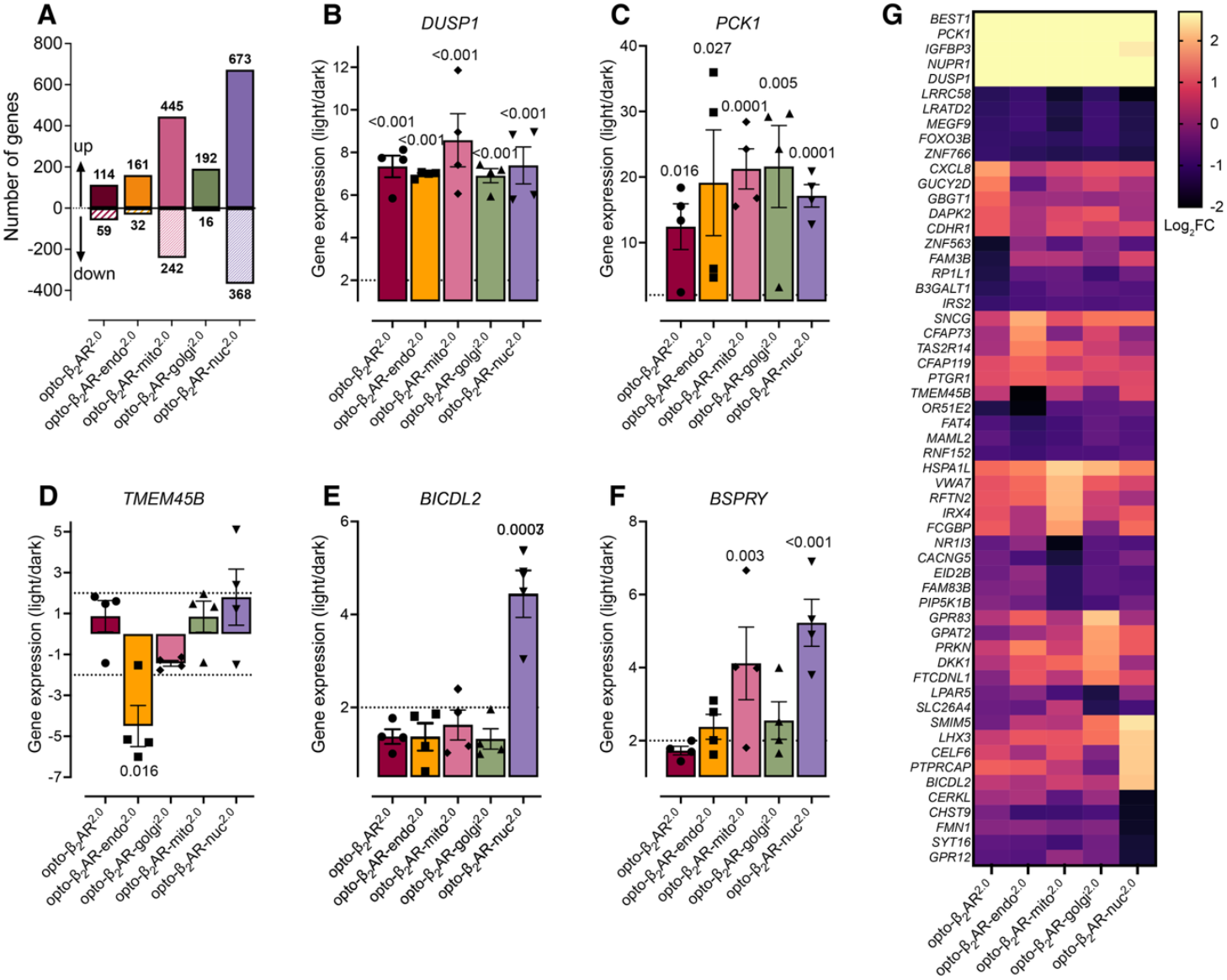
Targeted opto-β_2_AR^2.0^s induce unique gene transcriptional profiles in response to light. RNA-seq was performed following light stimulation of HEK293 cells stably transfected with the opto-β_2_AR^2.0^, opto-β_2_AR^2.0^-endo, opto-β_2_AR^2.0^-golgi, opto-β_2_AR^2.0^-mito, or opto-β_2_AR^2.0^-nuc (n=4). **A**) The number of genes (indicated above/below the bars) that were significantly altered in response to light activation of the untargeted or targeted opto-β_2_AR^2.0^s (>2-fold change, adjusted p-value <0.05). Light-induced significant changes in the expression of **B**) *DUSP1*, **C**) *PCK1*, **D**) *TMEM45B*, **E**) *BICDL2*, and **F**) *BSPRY* (n=4). Symbols show signed fold-change values derived from the log_2_(fold change) from independent experiments (retaining both the strength and direction of gene expression changes), bars show mean and error bars show SEM. Dotted lines indicate the 2-fold threshold for a change in gene expression, with adjusted p-values shown for significantly altered genes. **G**) Heat map of the top five up- and down-regulated protein coding genes that were commonly observed, or unique to cells stably transfected with the opto-β_2_AR^2.0^, opto-β_2_AR^2.0^-endo, opto-β_2_AR^2.0^-golgi, opto-β_2_AR^2.0^-mito, or opto-β_2_AR^2.0^-nuc (n=4). Only two genes were uniquely downregulated by the opto-β_2_AR^2.0^-golgi. Data shows average log_2_(fold change).

Light activation of the untargeted opto-β_2_AR^2.0^ altered the expression of 173 genes (>2-fold change and adjusted p-value <0.05), with 114 upregulated and 59 downregulated (Figure 6A, Supplementary Figure 8A, Supplementary Table 3). Several of these genes, including *DUSP1* and *PCK1*, are also regulated by ligand activation of the wild-type β_2_AR (*10, 52*), (Figure 6B-C, Supplementary Figure 8F). These genes formed part of a group of 72 genes that were commonly regulated by light activation of the opto-β_2_AR^2.0^ across all intracellular locations (Supplementary Figure 8F-G, Supplementary Table 4).

The light-activated opto-β_2_AR^2.0^-endo regulated the expression of 193 genes, including 29 genes (e.g. *TMEM45B*) that were uniquely regulated by opto-β_2_AR^2.0^-endo and not by any other opto-β_2_AR^2.0^ (Figure 6A, 6D, Supplementary Figure 8B, 8G). The genes regulated following stimulation of opto-β_2_AR^2.0^-endo showed a large degree of overlap with opto-β_2_AR^2.0^-nuc (145 shared genes, Supplementary Figure 8H), consistent with the established role of endosomal signalling proximal to the nucleus in gene regulation (*53*). Activation of the opto-β_2_AR^2.0^-nuc produced the largest transcriptional response (1041 genes), including 500 uniquely regulated genes such as *BICDL2*, not observed with any other opto-β_2_AR^2.0^ (Figure 6A, 6E and Supplementary Figure 8E, 8G). Across all locations, the largest overlap in commonly regulated genes occurred between opto-β_2_AR^2.0^-nuc and opto-β_2_AR^2.0^-mito (510 shared genes), including *BSPRY* (Figure 6F, Supplementary Figure 8H). Notably, the opto-β_2_AR^2.0^-mito induced the second-largest transcriptional response (687 genes), with 146 unique genes regulated at this location (Figure 6A, Supplementary Figure 8C, 8G). Finally, light activation of opto-β_2_AR^2.0^-golgi led to a similar number of transcriptional changes as activation of the untargeted opto-β_2_AR^2.0^, and opto-β_2_AR^2.0^-endo (208 genes), with 26 genes uniquely regulated at this location (Figure 6A, Supplementary Figure 8D, 8G).

To visualise the differences in the transcriptional profile across the targeted opto-β_2_AR^2.0^s we generated a heat map showing the top 5 upregulated and top 5 downregulated protein-coding genes in response to light activation (Figure 6G). This included genes commonly regulated across all receptors, as well as the top 5 uniquely up- and down-regulated genes for each location (except for opto-β_2_AR^2.0^-golgi where only two protein coding genes were uniquely downregulated; Supplementary Figure 8G). This analysis demonstrated that light stimulation induced distinct transcriptional profiles for each targeted opto-β_2_AR^2.0^ (Figure 6G).

## Discussion

In the present study, we adapted a chimeric receptor based on rhodopsin and the β_2_AR (opto-β_2_AR^2.0^) by adding C-terminal targeting sequences – this resulted in organelle-restricted expression of the opto-β_2_AR^2.0^ at early endosomes, the Golgi, mitochondria, or the nucleus. As the opto-β_2_AR^2.0^ is activated by light *in lieu* of ligand, this system allows us to compare the impact of activation of a receptor by the same stimulus at distinct locations in the cell. We found that the opto-β_2_AR^2.0^ could increase cAMP accumulation, ERK phosphorylation, and alter gene transcription at all intracellular organelles tested, with the respective magnitude, duration, and fingerprint of these signals dependent on receptor location.

Overall, our data provide orthogonal support that the competent platforms for intracellular GPCR signalling can be extended beyond the endosomal network to the membranes of the Golgi, the mitochondria and the nucleus. Whether these locations are ever sites of signalling for the wild-type β_2_AR remains to be determined, and the increasing availability of sensitive methods to detect endogenous β ARs (e.g. selective fluorescent agonists (*54*) and nanobodies (*9, 17*)) will likely facilitate such investigations. Previous studies have shown an increase in Gα_s_ recruitment to the Golgi in HEK293 cells following a prolonged stimulation with isoprenaline (*55*). Whether this is associated with the trafficking of the β_2_AR to this location is currently unknown. Beyond the β_2_AR, the Golgi is a well-recognised site for active GPCRs, that can either exist *in situ* (e.g. β_1_AR activation of cAMP and IP_3_ via an Epac/PLCε/mAKAPβ complex (*13, 14*)) or are trafficked from the plasma membrane following internalisation to activate a second phase of signalling (e.g. the thyroid stimulating hormone, TSH, receptor increases cAMP transiently at the plasma membrane, followed by sustained cAMP signalling from the trans-Golgi network (*12*)). Similarly, GPCRs have been identified *in situ* at the mitochondria (e.g. cannabinoid type 1, CB_1_, receptors decrease cAMP and PKA activity to influence mitochondrial respiration (*18*)). GPCRs can also either be trafficked to the nucleus (e.g. the proteinase-activated receptor, PAR2, changes its transcriptional effects as it traffics along microtubules from the plasma membrane to the nucleus (*56*)) or exist there *in situ* (e.g. metabotropic glutamate receptor, mGlu_5_, couples to Gα_q/11_ to increase calcium, calcium/calmodulin-dependent protein kinase IV, and activate the transcription factor Elk-1(*20, 21*)).

Although the opto-β_2_AR^2.0^ was able to activate both cAMP and ERK signalling at each intracellular organelle, we observed distinct differences in the temporal profiles of ERK phosphorylation at each location. The untargeted and mitochondrial opto-β_2_AR^2.0^ induced transient increases in ERK phosphorylation which returned to baseline by 30 min. In contrast, the endosomal, Golgi and nuclear opto-β_2_AR^2.0^ induced sustained increases in ERK phosphorylation at 30 min that were maintained for at least 4 h. Distinct temporal profiles of ERK activity have been reported for GPCRs that undergo internalisation to endosomes; activation of the neurokinin 1 receptor at the plasma membrane induces a transient increase in cytosolic ERK activity, the receptor is then internalised and can initiate sustained increases in ERK that propagate to the nucleus (*7, 8*). Similarly, activation of the wild-type β_2_AR leads to a transient increase in ERK activity when measured over the whole cell, however, this is sustained when measured directly at endosomes (*17*). Whether the initial transient signal is due to differential regulation of ERK signalling closer to the plasma membrane, or merely due to a time-dependent decrease in overall receptor activity at the plasma membrane due to receptor internalisation and trafficking, is unclear. As the targeted opto-β_2_AR^2.0^s are permanent resident proteins of the organelles, we can eliminate receptor trafficking as an explanation for transient signalling profiles. Our data instead suggests that there are fundamental, location-driven differences in the regulation of ERK signalling, such as differential compartmentalisation of phosphatases (*57*) or regulators of upstream components of this pathway, such as Ras (*58*). Future studies will focus on identifying the local interactome of the targeted opto-β_2_AR^2.0^s to mechanistically understand these differences.

The opto-β_2_AR^2.0^-endo, which exists within the endosomal network *in situ* and does not require β-arrestins to facilitate endocytosis from the plasma membrane, also allows us to dissociate the processes of receptor trafficking from endosomal G protein signalling. The ability of opto-β_2_AR^2.0^-endo to increase cAMP and ERK phosphorylation strongly implies that the signalling machinery is either present in endosomes or can be trafficked or recruited to endosomes *independently* of β-arrestin-dependent receptor endocytosis. Consistent with our observations, Gα_s_ can constitutively sample multiple intracellular membranes independently of receptor activation (*55*), and the direct activation of Gα_s_ (independently of GPCRs) can drive the relocation of plasma membrane-localised adenylyl cyclase 9 (AC9) to endosomes (*59*). Future studies should examine whether the activation of a GPCR that resides at the endosome (or other organelles) *in situ* results in the recruitment of regulatory proteins, such as GRKs and β-arrestin, that are characteristic events following activation of their plasma membrane counterparts.

While it is clear that intracellular organelles can host GPCRs and their signalling machinery, the exact location and orientation of the receptors (particularly in organelles with a dual membrane structure) and the proteins that comprise the local signalling cascades, are not well established. Based on the data presented here, we can start to build some hypotheses around this question. Visualisation of the mCherry-opto-β_2_AR^2.0^-nuc together with Venus-lamin B showed mCherry fluorescence immediately adjacent and to the outside of the inner nuclear membrane. Lamin B resides on the nucleoplasmic side of the inner nuclear membrane. This therefore suggests that the opto-β_2_AR^2.0^-nuc also resides on the inner nuclear membrane with the N-terminus facing the luminal cavity, and the C-terminus facing into the nucleus itself. This is entirely consistent with the topology of GPCRs during synthesis in the ER (*60*), and previous reports of the location and orientation of the α_1A_AR (*61*). The mitochondria is another organelle where the location and orientation of resident GPCRs is of great interest. Both the cannabinoid CB_1_ receptor (*18*) and the mouse melatonin MT_1_ receptor (*62, 63*) are thought to reside on the outer mitochondrial membrane with C-termini facing into the intermembrane space, and therefore would signal from the cytosol directly to the intermembrane space. Our imaging studies using three mitochondrial markers suggest that the opto-β_2_AR^2.0^-mito resides on the inner mitochondrial membrane. This location is consistent with reports of the location of the angiotensin AT_2_ receptor (*64*) and the serotonin 5-HT_4_ receptor (*65*). At this point, the orientation of GPCRs that reside on the inner mitochondrial membrane remains unclear, and how signalling either into the intermembrane space or the mitochondrial matrix occurs is an area of continuing research. For example, while the mitochondrial matrix contains proteins required for cAMP signalling and transcription, cAMP in this compartment appears to be dependent on the activation of soluble AC (*66*). Soluble AC is activated by calcium or bicarbonate (not G proteins) (*67*), suggesting that a GPCR positioned in the inner mitochondrial membrane may influence the transport of these ions into the matrix to increase cAMP. Future studies are required to elucidate the precise signalling pathways involved in the regulation of transcription by the opto-β_2_AR^2.0^ in different intracellular compartments.

In summary, we have engineered a synthetic system that allows investigation of the impact of receptor location on cell signalling. By using optogenetic chimeric receptors targeted to different intracellular membranes we can directly compare the influence of location alone on the dynamics of receptor signalling and transcriptional responses. In so doing, we have demonstrated that endosomes, the Golgi, mitochondria and the nucleus have access *in situ* to the cellular machinery required for activation of cAMP and ERK signalling, and transcriptional initiation. Our targeted optogenetic toolbox represents a valuable approach that can be used to understand the role of GPCR location in the coordination of cellular responses.

## Methods

### Reagents

9-cis-retinal, donkey anti-sheep IgG (H+L) cross-adsorbed secondary antibody Alexa Fluor™ 488 (A11015), goat anti-rabbit IgG (H+L) cross-adsorbed secondary antibody Alexa Fluor™ 488 (A11008), forskolin, 3-isobutyl-1-methyxanthine (IBMX), poly-D-lysine and polyethylenimine (PEI) were purchased from Sigma-Aldrich. CellLight™ Mitochondria-GFP BacMam 2.0 and Golgi-GFP BacMam 2.0, Dulbecco’s modified Eagle’s medium (DMEM) with high glucose, pyruvate and GlutaMAX, fetal bovine serum (FBS), Geneticin™ selective antibiotic (G418 sulfate), and Hoescht were from Thermo Fisher Scientific. Paraformaldehyde (16% v/v) was purchased from ProSciTech. Anti-EEA1 (C45B10) rabbit monoclonal antibody (3288T) was purchased from New England Biolabs. Sheep anti-TGN46 antibody (AHP500GT) was purchased from BioRad. Rabbit anti-TOMM20 [EPR15581-54] monoclonal antibody (ab232589) was from Abcam.

### cDNAs

Plasmids encoding opto-β_2_AR^2.0^ (*27*), and Rab5a-Venus (*7*) were previously described. KRas-Venus, Venus-PTP1b, and Venus-giantin were a gift from N. Lambert (*68*). nucEKAR cerulean/venus (Addgene plasmid 18681) was a gift from K. Svoboda (*51*), Venus-mito-7 (Addgene plasmid 56619) and Venus-Lamin B (Addgene plasmid 56616) were a gift from M. Davidson.

mCherry-opto-β_2_AR^2.0^-MITO2 was synthesised and sub-cloned into pTwist-EF1α by Twist Biosciences.

Indicated targeting sequences (Table 1) were ordered as either double-stranded gBlock gene fragments (sequences EE2, ER2, ER3, GOL2, GOL3, NUC1, and NUC2) or single-stranded complementary oligonucleotides (sequences EE1, ER1, GOL1, and MITO1) (Integrated DNA Technologies). In each case, targeting sequences were cloned onto the C-terminus of the opto-β_2_AR^2.0^ (pcDNA3.1-, Thermo Fisher Scientific) using Golden Gate Assembly with each targeting sequence followed by a stop codon. A fluorescent mCherry tag was cloned onto the N-terminal end of each protein to enable visualisation. Construct sequences were verified at MicroMon Next-Generation Sequencing (Monash University, Australia).

### Cell culture

HEK293 cells (American Type Culture Collection; negative for mycoplasma contamination) were cultured in DMEM supplemented with 5% (v/v) FBS. Cells were transiently transfected with linear PEI at a DNA:PEI ratio of 1:6 (*69*), and all assay plates were pre-coated poly-D-lysine (5 μg/cm^2^).

HEK293 cells stably transfected with opto-β_2_AR^2.0^, opto-β_2_AR^2.0^-endo, opto-β_2_AR^2.0^-mito, opto-β_2_AR^2.0^-golgi or opto-β_2_AR^2.0^-nuc were generated for RNA-seq experiments. Cells in T25 flasks were transfected with 550 ng untargeted or targeted opto-β_2_AR^2.0^ using PEI. After 48 h, media was replaced with fresh culture media supplemented with 600 μg/mL G418 sulfate. Media was replaced every 2-3 days and cells were expanded as required. After 2 weeks of selection, cells were sorted for mCherry fluorescence by fluorescence activated cell sorting (Sandy Fung, Monash Institute of Pharmaceutical Sciences Imaging, FACS, and Analysis Core) under sterile conditions using a Beckman Coulter MoFlo Astrios, with untransfected HEK293 cells as a negative control. Sorted cells were expanded to generate polyclonal stable cell lines. Stable cell lines were placed into growth media containing 600 μg/mL G418 sulfate every 2 weeks to maintain selection pressure.

### Light activation of optogenetic receptors

For confocal imaging of receptor location, cells were illuminated with the OPSL 488 laser (493-582 nm) of a Leica TCS SP8 confocal microscope. For cAMP assays, the Xenon light source (460-540 nm using the 500-80 excitation filter) of the CLARIOstar multimode microplate reader (BMG Labtech) was used to illuminate cells with 1-1000 flashes of light. For ERK phosphorylation assays and RNA-seq experiments, cells were illuminated with our previously described modular LED shelving system (*29*). Briefly, the apparatus consists of a 5-drawer shelving unit with acrylic sheets custom cut to accommodate assay plates and fits easily in a standard cell culture incubator. A green LED array (536-596 nm; used at 32.49±3.42 μW/cm^2^) was placed on the lower shelf (drawer slide 1) to illuminate cells in clear-bottom assay plates on a shelf positioned two drawer slides above (drawer slide 3; Figure 1B). Assay plates were covered with a black opaque lid to avoid incidental or reflected light entering from the top or sides of the plate. A small portable fan was placed to the side of the LEDs to facilitate the distribution of heat generated by the LED array (Figure 1B).

### Confocal imaging

HEK293 cells were co-transfected in 96-well plates (4×10^4^ cells per well) with untargeted or targeted opto-β_2_AR^2.0^ (75 ng/well) and an intracellular marker (25 ng/well). CellLight mitochondria-GFP or CellLight Golgi-GFP were added to cells 24 h after transfection. Forty-eight h post transfection, live cells were imaged in culture media. For nuclear Hoescht staining, media was removed and cells were incubated with Hoescht in PBS for 15 min at 37°C. To visualise the inactive, untargeted opto-β_2_AR^2.0^, cells were fixed in the dark by washing three times in ice-cold PBS, prior to fixation in 4% (v/v) paraformaldehyde for 15 min at room temperature. Cells were thrice-washed with PBS and remained in PBS until imaging.

For early endosome immunostaining with anti-EEA1 antibody, fixed cells were blocked for 1 h at room temperature in blocking buffer (5% v/v goat serum, 0.3% v/v Triton-X 100 in PBS). Cells were then incubated with anti-EEA1 antibody (1:200 in dilution buffer; 1% w/v BSA, 0.3% v/v Triton-X 100 in PBS) for 1 h at room temperature, thrice washed with PBS for 5 min, followed by a 1 h incubation with goat anti-rabbit AlexaFluor-488 secondary antibody (1:1000 in dilution buffer) also at room temperature. For immunostaining of the trans-Golgi network with anti-TGN46 antibody, fixed cells were blocked overnight at 4°C in blocking buffer (5% v/v horse serum, 0.3% v/v Triton-X 100 in PBS). Cells were then incubated for 1 h at room temperature with anti-TGN46 antibody (1:500 in dilution buffer), thrice washed with PBS for 5 min, followed by a 1 h incubation at room temperature with donkey anti-sheep AlexaFluor-488 secondary antibody (1:2000 in dilution buffer). For immunostaining of the outer mitochondrial membrane with anti-Tom20 antibody, fixed cells were blocked for 1 h at room temperature in blocking buffer (5% v/v goat serum, 0.3% v/v Triton-X 100 in PBS). Cells were then incubated with anti-Tom20 antibody (1:1000 in dilution buffer) for 1 h at room temperature, thrice washed with PBS for 5 min, followed by a 1 h incubation with goat anti-rabbit AlexaFluor-488 secondary antibody (1:1000 in dilution buffer) also at room temperature. Cells were thrice washed (5 min each) in PBS and imaged in PBS.

Imaging was performed using a Leica TCS SP8 confocal microscope equipped with a dry immersion 20X or 40X 0.75 NA HC PL APO CS2 objective. Diode 405 (410-483 nm emission filter), OPSL 552 (582-730 nm) and OPSL 488 (493-582 nm) lasers were used to image Hoescht, mCherry and Venus/AlexaFluor-488/CellLight-GFP fluorescence, respectively. Sequential scanning was used to minimise crosstalk between fluorophore channels. All transfection conditions were performed in duplicate and two fields of view were captured per well.

Data were analysed with the FIJI distribution of ImageJ (*70*), using a custom analysis pipeline (https://bridges.monash.edu/; doi 10.26180/24770562) and a researcher who was blinded. Cells with both the organelle marker and mCherry-tagged opto-β_2_AR^2.0^ were selected for analysis using the “Mark Out Cells” script. A region lacking cells was selected to define the background. The amount of mCherry fluorescence from the opto-β_2_AR^2.0^ that was found in the same pixels identified by the marker was calculated using the “Analyse Cells” script. The script creates a mask using the marker image. The raw integrated density of the mCherry-opto-β_2_AR^2.0^ in the entire image compared to that in the mask area is calculated. Cells were considered for analysis if the signal for the marker was more than three times the background signal.

As this quantification could be influenced by the relative expression level of each of the proteins (protein of interest, and location marker) or the efficiency of the stain, we determined the upper and lower thresholds of this co-incident analysis. As a positive control, we quantified the amount of fluorescence from a nuclear-localised ERK biosensor (nucEKAR (*51*)) that occurred within pixels defined by the DNA-binding Hoescht stain (Supplementary Figure 1A-B). As a negative control, we quantified the amount of fluorescence from the KRas-Venus plasma membrane marker that occurred within pixels defined by the DNA-binding Hoescht stain (Supplementary Figure 1A, 1C). We determined the median percentage of co-incident localisation from all cells that expressed the Venus-tagged markers within an image and averaged this value from three independent biological experiments (Supplementary Figure 1B-D). This allowed us to define the upper and lower limits of our quantification system, as 89.9% and 24.2%, respectively (Supplementary Figure 1D). Values from all analysed cells from three independent experiments were plotted using GraphPad Prism, with values normalised to the defined negative (KRas-Venus plasma membrane marker vs Hoescht DNA stain) and positive (nucEKAR vs Hoescht DNA stain) controls. Data are shown as box plots, defining the median and interquartile range.

For line scan analysis of representative images, the cell was outlined using a merged image (including brightfield) and the centre of the cell was determined using the centroid measurement in ImageJ. A straight line was drawn through the centroid, across a region of a cell containing both mCherry and marker fluorescence. If this central line did not include much of the organelle marker, a second line was drawn through an area of high marker fluorescence. The ‘plot profile’ function was used to quantify the intensity of pixels in both mCherry and marker channels. Data were normalised to the maximal intensity for each channel and plotted using GraphPad Prism.

### Receptor expression

Receptor expression was determined by measuring mCherry fluorescence using the CLARIOstar multimode plate reader (excitation and emission filters 565-25 and 635-50 nm, respectively). Cells were illuminated with 20 flashes of light for a run time of 16 seconds. Data was collected using the MARS Data Analysis Software.

### cAMP accumulation assays

HEK293 cells were transfected with 550 ng untargeted or targeted opto-β_2_AR^2.0^ in 6-well plates (9×10^5^ per well). After 48 h, accumulation of intracellular cAMP was quantified in triplicate using a CisBio Gs dynamic cAMP kit, following the manufacturer’s instructions. Cells were incubated at 37°C in the dark with 9-cis-retinal (10 μM) for 1 h prior to light activation. Under red light (to avoid activation of light-responsive receptors), cells were transferred to a 384-well plate at 5,000 cells per well in the presence of 10 μM 9-cis-retinal and 500 μM IBMX in stimulation buffer. “Dark” control cells did not receive light stimulation, and 100 μM forskolin was used as a positive control. After light stimulation with the CLARIOstar, cells were placed on ice. Detection reagents (cAMP-d2 antibody and anti-cAMP-cryptate antibody) were separately prepared in lysis buffer and 5 μL of each was added to all wells. Plates were then sealed, covered in foil and left at room temperature for 1 h. Plates were read using a PerkinElmer Envision multi-mode plate reader (excitation 665 nm, emission 620 nm). Data were analysed against a standard curve, and light-response curves were fit using non-linear regression and the log(agonist) vs. response (three parameter) equation in GraphPad Prism.

### ERK phosphorylation assays

HEK293 cells were transfected with 5 μg untargeted or targeted opto-β_2_AR^2.0^ in 10 cm dishes (4×10^6^ cells per dish). After 24 h, cells were transferred to black, clear-bottom 96-well ViewPlates (2.5×10^4^ cells/well) and allowed to adhere for 6 h. Culture media was then replaced with serum-free media supplemented with 10 μM 9-cis-retinal for 18 h. For time course experiments, black stickers were placed on the clear bottom of the assay plate with a black lid, and removed at specific time intervals to allow cells to be exposed to light. “Dark” control cells did not receive light stimulation (i.e. black stickers were not removed), and a 10 min incubation with 10% (v/v) FBS was used as a positive control. ERK1/2 phosphorylation was quantified in duplicate using the CisBio phospho-ERK1/2 (Thr202/Tyr204) kit, according to the manufacturer’s instructions. After light stimulation, supplemented lysis buffer (including blocking reagent) was added to the plate and allowed to shake for 30 min. Cells were then lysed overnight at -20°C. The next day, 16 μL of cell lysate was transferred to a 384-well white ProxiPlate to which 2 μL of each detection antibody (phospho-ERK1/2 d2 antibody and phospho-ERK1/2 Eu cryptate antibody prepared in detection buffer) were added. Plates were read using a PerkinElmer EnVision multi-mode plate reader (excitation 665 nm, emission 620 nm). Given the variability of transient transfection, filtering steps were applied to account for biological replicates with very low transfection efficiency. Biological replicates were excluded if (1) the light-stimulated ERK phosphorylation response was less than 10% of the dark response or (2) light induced a decrease in ERK phosphorylation of more than 15% compared to the dark response. The “peak” response was determined as the time point that induced the largest increase in ERK phosphorylation. Data are expressed as either fold change compared to dark (30 min time courses; Figure 1F and Figure 5A, C, E, G), relative fluorescence units (statistical analysis of 2 min and 30 min responses; Figure 1G and Figure 5B, D, F, H) or fold change relative to peak (4 h time courses; Figure 5I-M) to allow appropriate statistical analyses.

### RNA sequencing

HEK293 cells stably transfected with untargeted or targeted opto-β_2_AR^2.0^ were seeded in 6-well plates (7×10^5^ cells/well) in complete culture medium and allowed to adhere for 6 h. Culture media was then replaced with serum-free media supplemented with 10 μM 9-cis-retinal for 18 h. On the day of the experiment, cells were stimulated with light for 2 h. Control (“dark”) plates did not receive 9-cis-retinal incubation nor light stimulation (plates were wrapped in foil and placed on the same shelf in the cell culture incubator). RNA was extracted from cells under red (non-activating) light using the RNeasy Mini kit (Qiagen). Cell treatment and RNA extraction were performed on four separate occasions. Transcriptomic sequencing was performed by the Australian Genome Research Facility (AGRF).

Raw sequencing reads were assessed for quality using FastQC (v0.11.7; https://www.bioinformatics.babraham.ac.uk/projects/fastqc/), and adapter trimming was performed using Trimmomatic (v0.39) (*71*). High-quality reads were aligned to the human ensemble GRCh38 genome using HISTAT2 (v2.2.1) (*72*), and the data were uploaded to the Galaxy web platform (*73*) using the UseGalaxy Australia public server (https://usegalaxy.org.au) for further analysis. Gene-level counts were quantified using featureCounts (*74*) (Galaxy version 2.1.1+galaxy0), and differential expression analysis was conducted using the limma-voom (*75*) pipeline (Galaxy version 3.58.1+galaxy0). Genes with low expression (<0.03125 CPM, equivalent to log_2_CPM < -5) across all samples were excluded from further analysis. Due to the addition of 9-cis-retinal as a chromophore in our light-treated samples, we also removed all genes associated with retinal signalling and metabolism, as found in the KEGG “Retinal metabolism”, Gene Ontology Biological Process “Retinoic acid receptor signalling pathway”, and Reactome “Signaling by retinoic acid” gene lists (accessed January 2026; 96 genes total). Genes were considered differentially expressed if the adjusted p-value was < 0.05 and the absolute log_2_ fold change exceeded 1 (equivalent to a 2-fold change). Protein coding genes were identified using Ensembl BioMart. Venn diagrams were created using the Compare Lists (Multiple List Comparator) tool by molbioltools (https://molbiotools.com/listcompare.php). All other plots and heatmaps were created using GraphPad Prism.

### Statistics and reproducibility

Statistical analyses were performed using GraphPad Prism software (v10.6.1), and all relevant information is included in the figure legends (Figures 1–6, Supplementary Figures 1–8, Supplementary Tables 1-4). Data are expressed as mean ± standard error of the mean (SEM) unless otherwise indicated, with sample sizes indicated in the figure legends. Normal distribution of data was confirmed using QQ plots in GraphPad Prism; p<0.05 was considered statistically significant.

## Data Availability Statement

The RNA sequencing data generated in this study have been deposited in the Gene Expression Omnibus (GEO) under accession number [TBC]. All other data supporting the findings of this study are available within the paper and its Supplementary Information.

## Acknowledgements

We thank members of the Spatial Organisation of Signalling Laboratory for valuable discussions of the project. The authors acknowledge the use of instruments at the Monash Institute of Pharmaceutical Sciences (MIPS) Imaging, FACS and Analysis Core.

M.L.H. is a Viertel Senior Medical Research Fellow supported by The Cross Family and The Frank Alexander Charitable Trusts. A.M.E. is a Victorian Department of Health and Human Services Victorian Cancer Agency Mid-Career Fellow (MCRF21036). This research was supported by a Monash University Joint Medicine Pharmacy grant to M.L.H. and H.J.

## Author Contributions

M.L.H. and H.J. conceived the project

M.L.H. supervised the project and designed experiments

C.M. designed experiments, performed all imaging, cAMP and ERK experiments, and analysed all data

B.L. performed sample preparation for RNA-seq

C.M. and M.L.H. interpreted results

C.G.G. made LED arrays and shelves, supervised the set-up of the optical shelving system

A.M.T., C.M., H.J. and A.M.E. designed and engineered targeted receptors

C.N. wrote and developed semi-automated image analysis protocols

C.M., M.L.H. and A.M.E. wrote the manuscript

All authors edited the manuscript

