## Supplemental Material for "Intracellular targeting of a light-activated GPCR elicits spatially selective signalling"

**Supplementary Information**

Supplementary Table 1
Supplementary Tables 2-4 (excel files)
Supplementary Figures 1-9

**Supplementary Table 1. C-terminal organelle targeting sequences.** Complete list of all identified C-terminal sequences. The table shows if they were tested in this study, and if so, whether they conferred correct localisation of the opto-$\beta_2$AR$^{2.0}$.

| Target Organelle | Name | Sequence | Protein Source | Tested? | Correct localisation of opto-$\beta_2$AR$^{2.0}$? *Reference figures* |
|---|---|---|---|---|---|
| Early endosomes | EE1 | FDNPVY | Low density lipoprotein receptor (*33*) | Yes | No<br>*Supplementary Figure 2A-C, 2E* |
| | EE2 | IEGLYDMHIQLQSVPFLHWEAPVTSHSLTAREVMSTPVTCLRRREKVGVIVDVLSDTASNHNGFPVVEHADDTQPARLQGLILRSQLIVLLKHKVFVERSNLGLVQRRLRLKDFRDAYPRFPPIQSIHVSQDERECTMDLSEFMNPSPYTVPQEASLPRVFKLFRALGLRHLVVVDNRNQVVGLVTRKDLARYRLGKRGLEELSLAQT | CLC-7 (*34*) | Yes | Yes<br>*Figure 2A-B*<br>*Supplementary Figure 2A, 2D-G* |
| Endoplasmic Reticulum | ER1 | RMRTR | RxRxR motif from $GABA_{B1}$ (*35*) | Yes | No<br>*Supplementary Figure 6A-C, 6F* |
| | ER2 | LLQKFQKDLNTGVFNNQENEILKQIVKHDREMVQAIAPINYPQMTTLNSTSSTTTPTSRMRTQ | HCN1 channel (*36*) | Yes | No<br>*Supplementary Figure 6A-B, 6D, 6F* |
| | ER3 | PVTCAAAAGGMGVGVGVGGSGFRNVYAEMLHFQSMCSCLW | Rat TASK1 (*37*) | Yes | No<br>*Supplementary Figure 6A-B, 6E-F* |
| | ER4 | TRRRRRSLSNVN | Kir6.2 (*37*) | No | |
| | ER5 | RNVYAEMLHFQSMCSCL | Rat TASK1 (*37*); truncated version of ER3 | No | |
| | ER6 | KYKSRRSFIDEKKMP | Adenovirus E19 *(76)* | No | |
| Golgi | GOL1 | FLSSR | F/L-L/I/V-X-X-R/K motif in glycosyltransferases from Yeast (*38*) | Yes | No<br>*Supplementary Figure 4A-C, 4F* |
| | GOL2 | RLLDSSNATWQRDQPDTHRLSRLDAHVMSM | Uukuniemi virus G1 membrane glycoprotein (*39*) | Yes | Yes<br>*Figure 3A-B*<br>*Supplementary Figure 4A, 4D, 4F-H* |
| *Trans-Golgi network* | GOL3 | HNKRKIIAFALEGKRSKVTRRPKASDYQRLNLKL | TGN38 (*40*) | Yes | No<br>*Supplementary Figure 4A-B, 4E-F* |
| Mitochondria | MITO1 | DRKRRSDPNFK | Tom20 (*41*) | Yes | Yes<br>*Figure 2C-D*<br>*Supplementary Figure 3A, 3C, 3E-G* |
| | MITO2 | DRKRRSD DRKRRSD | Tandem truncated version of MITO1 (*41*) | Yes | No<br>*Supplementary Figure 3A-B, 3D-E* |
| Nucleus | NUC1 | LKEKKKRTVAEEDQLHLDGQENKRRRHDSS | Rat pom121 (*42*) | Yes | Yes<br>*Figure 3C-D*<br>*Supplementary Figure 5A, 5C, 5E-F* |
| | NUC2 | HREARDEYHKKKYGVAWEKYCQRVPYRIFPYIY | Lamin B receptor (*43*) | Yes | No<br>*Supplementary Figure 5A-B, 5D-E* |
| | NUC3 | DRLRHR | Cytomegalovirus glycoprotein B *(77)* | No | |
| | NUC4 | SNSNNKNDANKNN [needs to be immediately after final helix] | baculovirus “bait” sequence *(78)* | No | |
| | NUC5 | KKFKR | AT1 receptor *(79)* | No | |

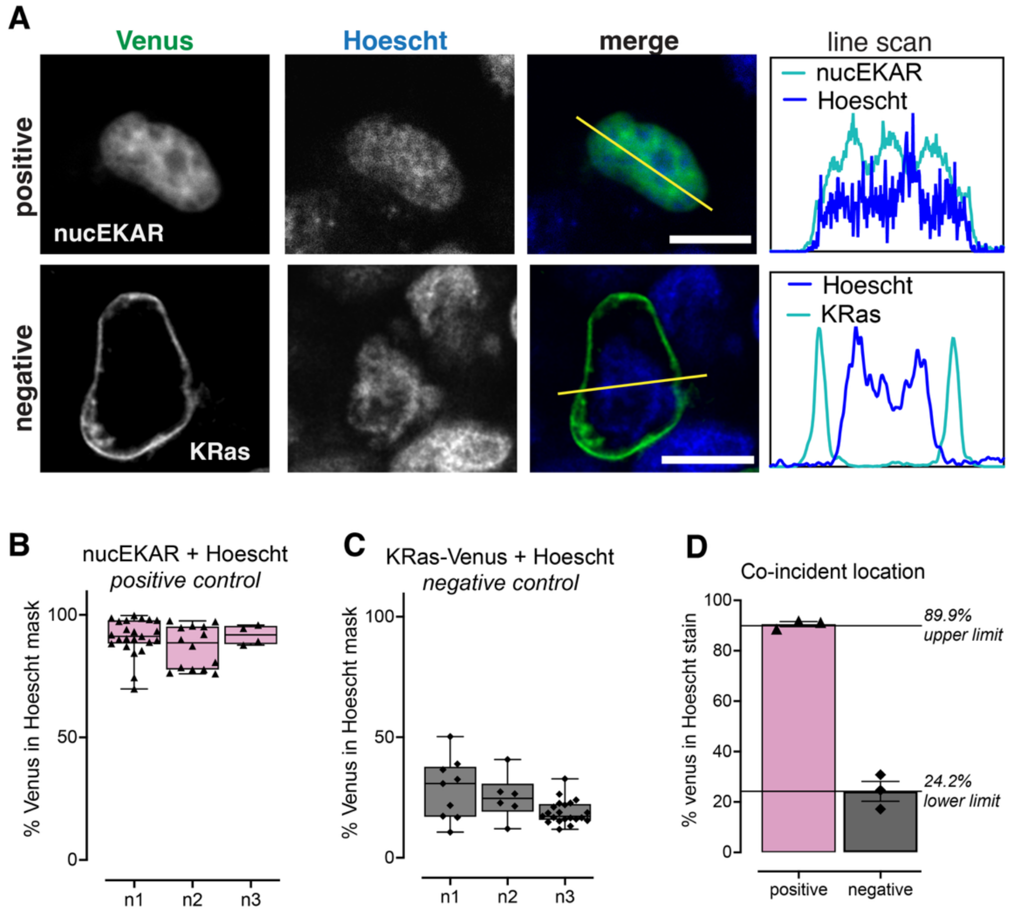


**Supplementary Figure 1. Establishing the upper and lower limits for semi-automated quantification of sub-cellular targeting.** HEK293 cells were transfected with a nuclear localised ERK biosensor (nucEKAR; positive control) or a plasma membrane marker (KRas-Venus; negative control) and nuclei were defined using a Hoescht stain. **A**) representative images of cells expressing the positive or negative control (left panel), Hoescht stain (middle panel) and merged image (right panel), with the yellow line indicating the region highlighted in the line scan intensity graph (far right panel). Scale bar is 10 μm. Quantification of the percentage of Venus fluorescence identified in the pixels defined by the Hoescht nuclear stain for the **B**) positive or **C**) negative controls (n=3). Symbols indicate individual cells, bar graphs show the median and minimum/maximum values. **D**) Averaged median co-incident location defining 89.9% as the upper limit, and 24.2% as the lower limit for the system. The quantification of opto-$\beta_2$AR$^{2.0}$ targeting (Figures 2 and 3, Supplementary Figures 2-6) was normalised to these upper and lower limits.

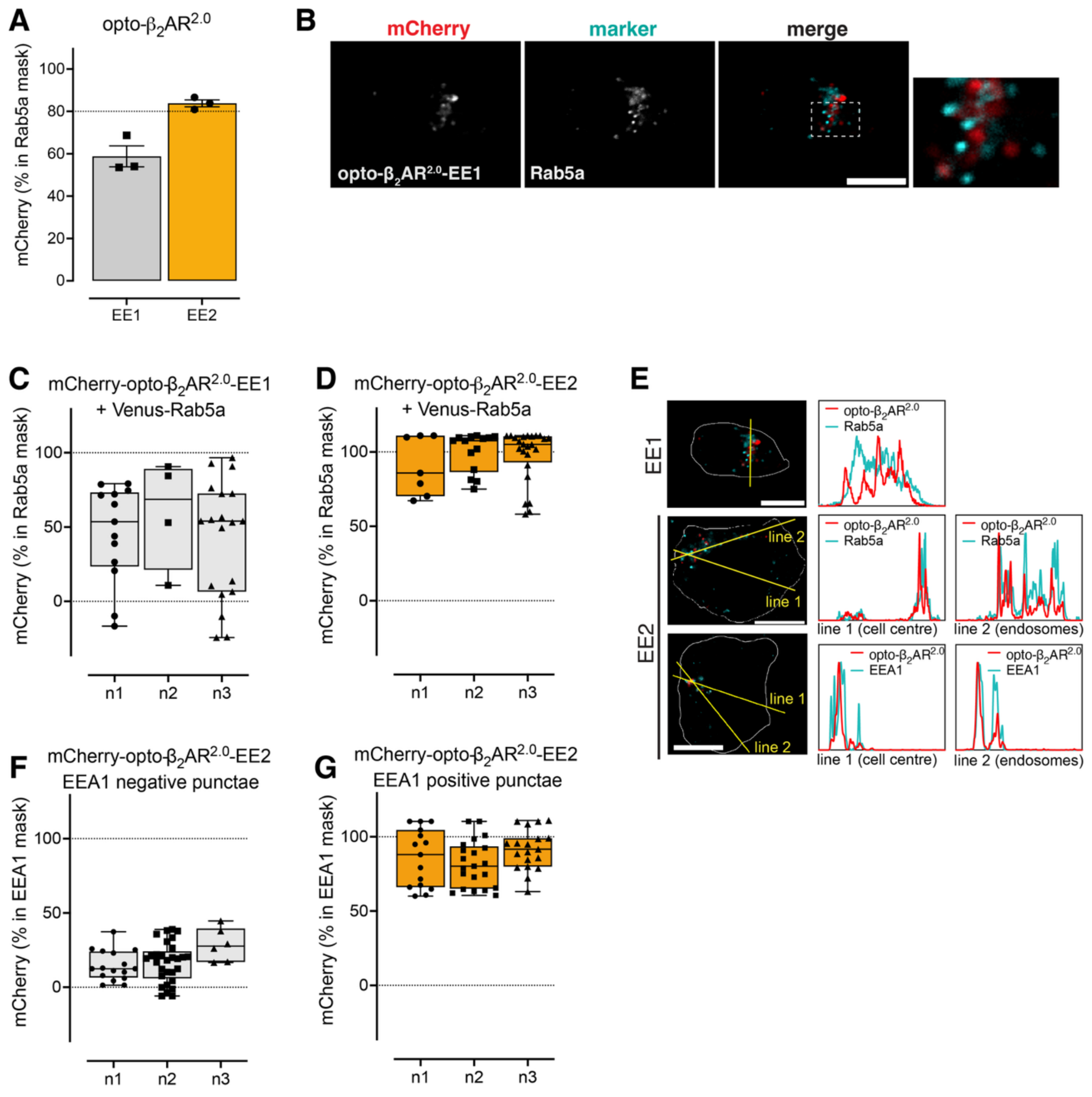


**Supplementary Figure 2. C-terminal sequence EE2, but not EE1, targeted the opto-$\beta_2$AR$^{2.0}$ to early endosomes.** HEK293 cells were transfected with the targeted mCherry-opto-$\beta_2$AR$^{2.0}$ and either co-transfected with a Venus-tagged intracellular marker, infected with a GFP-tagged CellLight marker, or immunostained with an antibody against an organelle resident protein. **A**) An initial screen compared the amount of mCherry fluorescence within the pixel area defined by Venus-Rab5a following C-terminal fusion of EE1 or EE2 sequences to mCherry-opto-$\beta_2$AR$^{2.0}$. Symbols show median values from all cells from each biological replicate (shown in C and D) the bar shows the mean and error bars show SEM (n=3). **B**) Representative images show the EE1 sequence does not localise the mCherry-opto-$\beta_2$AR$^{2.0}$ (left panel) to Venus-Rab5a (middle panel). Right panel shows merged image, and the inset (far right) shows the region boxed in white. Scale bar is 10 $\mu$m. **C-D**) The amount of mCherry-opto-$\beta_2$AR$^{2.0}$ fluorescence within the pixel area defined by the Venus-Rab5a marker was calculated for each cell in the image, following C-terminal fusion of the EE1 (C) or EE2 (D) sequences. Symbols show data from individual cells, bars show median and error bars show the minimum/maximum values (n=3). **E**) Line scan analysis of the representative merged images of the mCherry-opto-$\beta_2$AR$^{2.0}$ and organelle markers as displayed in B (this figure) and Figure 2B. The cell outline (white line, defined by brightfield image) was used to calculate the centroid, through which the yellow line was drawn for line scan intensity analysis (inset graphs). When this line did not sufficiently capture the organelle marker, a second line (line 2) was drawn through a region of high intensity of the marker by the blinded investigator. Scale bar is 10 $\mu$m. **F-G**) The amount of mCherry-opto-$\beta_2$AR$^{2.0}$-EE2 fluorescence within the pixel area defined by EEA1 immunostaining, with "EEA1 negative" (F) and "EEA1 positive" (G) refering to mCherry punctae that were negative or positive for EEA1 immunostaining, respectively. Symbols show data from individual cells, bars show median and error bars show the minimum/maximum values (n=3).

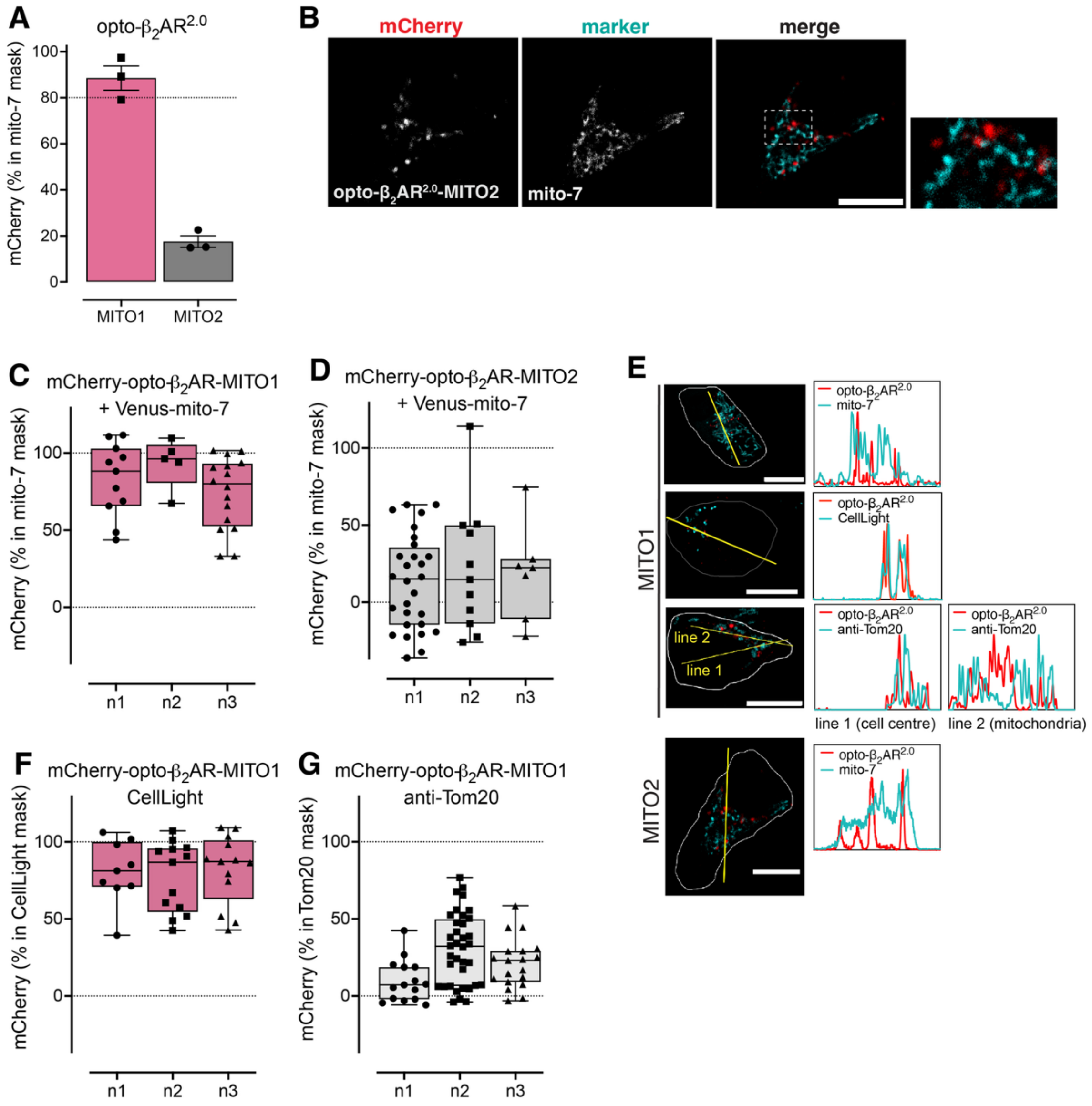


**Supplementary Figure 3. C-terminal sequence MITO1, but not MITO2, targeted the opto-$\beta_2$AR$^{2.0}$ to mitochondria.** HEK293 cells were transfected with the targeted mCherry-opto-$\beta_2$AR$^{2.0}$ and either co-transfected with a Venus-tagged intracellular marker, infected with a GFP-tagged CellLight marker, or immunostained with an antibody against an organelle resident protein. **A**) An initial screen compared the amount of mCherry fluorescence within the pixel area defined by Venus-mito7 following C-terminal fusion of MITO1 or MITO2 sequences to mCherry-opto-$\beta_2$AR$^{2.0}$. Symbols show median values from all cells from each biological replicate (shown in C and D) the bar shows the mean and error bars show SEM (n=3). **B**) Representative images show the MITO2 sequence does not localise the mCherry-opto-$\beta_2$AR$^{2.0}$ (left panel) to Venus-mito7 (middle panel). Right panel shows merged image, and the inset (far right) shows the region boxed in white. Scale bar is 10 $\mu$m. **C-D**) The amount of mCherry-opto-$\beta_2$AR$^{2.0}$ fluorescence within the pixel area defined by the Venus-mito7 marker was calculated for each cell in the image, following C-terminal fusion of the MITO1 (C) or MITO2 (D) sequences. Symbols show data from individual cells, bars show median and error bars show the minimum/maximum values (n=3). **E**) Line scan analysis of the representative merged images of the mCherry-opto-$\beta_2$AR$^{2.0}$ and organelle markers as displayed in B (this figure) and Figure 2D. The cell outline (white line, defined by brightfield image) was used to calculate the centroid, through which the yellow line was drawn for line scan intensity analysis (inset graphs). When this line did not sufficiently capture the organelle marker, a second line (line 2) was drawn through a region of high intensity of the marker by the blinded investigator. Scale bar is 10 $\mu$m. **F-G**) The amount of mCherry-opto-$\beta_2$AR$^{2.0}$-MITO1 fluorescence within the pixel area defined by the Mitochondria-GFP CellLight marker (F) or Tom20 immunostaining (G). Symbols show data from individual cells, bars show median and error bars show the minimum/maximum values (n=3).

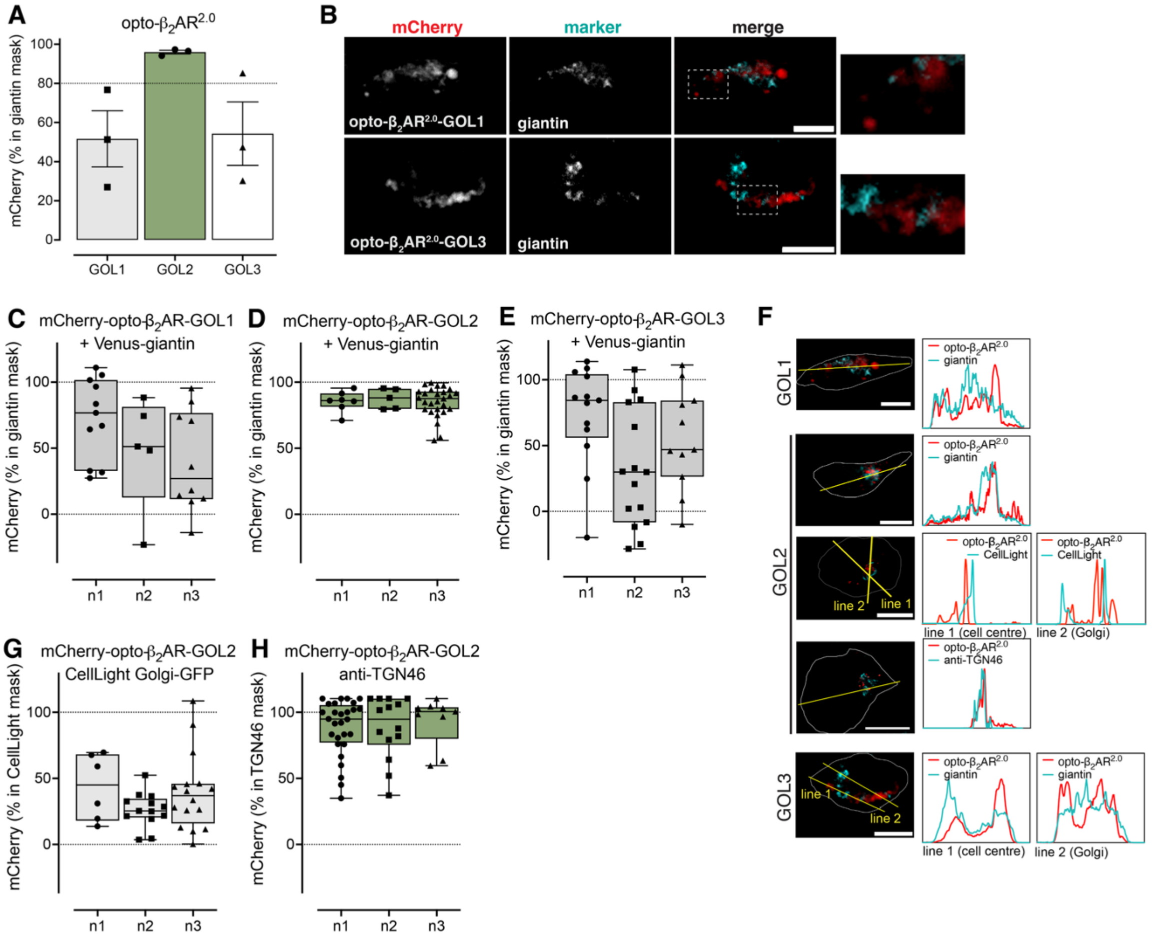


**Supplementary Figure 4. C-terminal sequence GOL2, but not GOL1 or GOL3, targeted the opto-$\beta_2AR^{2.0}$ to the Golgi.** HEK293 cells were transfected with the targeted mCherry-opto-$\beta_2AR^{2.0}$ and either co-transfected with a Venus-tagged intracellular marker, infected with a GFP-tagged CellLight marker, or immunostained with an antibody against an organelle resident protein. **A**) An initial screen compared the amount of mCherry fluorescence within the pixel area defined by Venus-giantin following C-terminal fusion of GOL1, GOL2 or GOL3 sequences to mCherry-opto-$\beta_2AR^{2.0}$. Symbols show median values from all cells from each biological replicate (shown in C-E), the bar shows the mean and error bars show SEM (n=3). **B**) Representative images show the GOL1 and GOL2 sequences do not localise the mCherry-opto-$\beta_2AR^{2.0}$ (left panel) to Venus-giantin (middle panel). Right panel shows merged image, and the inset (far right) shows the region boxed in white. Scale bar is 10 $\mu$m. **C-E**) The amount of mCherry-opto-$\beta_2AR^{2.0}$ fluorescence within the pixel area defined by the Venus-giantin marker was calculated for each cell in the image, following C-terminal fusion of the GOL1 (C), GOL2 (D), or GOL3 (E) sequences. Symbols show data from individual cells, bars show median and error bars show the minimum/maximum values (n=3). **F**) Line scan analysis of the representative merged images of the mCherry-opto-$\beta_2AR^{2.0}$ and organelle markers as displayed in B (this figure) and Figure 3B. The cell outline (white line, defined by brightfield image) was used to calculate the centroid, through which the yellow line was drawn for line scan intensity analysis (inset graphs). When this line did not sufficiently capture the organelle marker, a second line (line 2) was drawn through a region of high intensity of the marker by the blinded investigator. Scale bar is 10 $\mu$m. **G-H**) The amount of mCherry-opto-$\beta_2AR^{2.0}$-GOL2 fluorescence within the pixel area defined by the Golgi-GFP CellLight marker (G) or TGN46 immunostaining (H). Symbols show data from individual cells, bars show median and error bars show the minimum/maximum values (n=3).

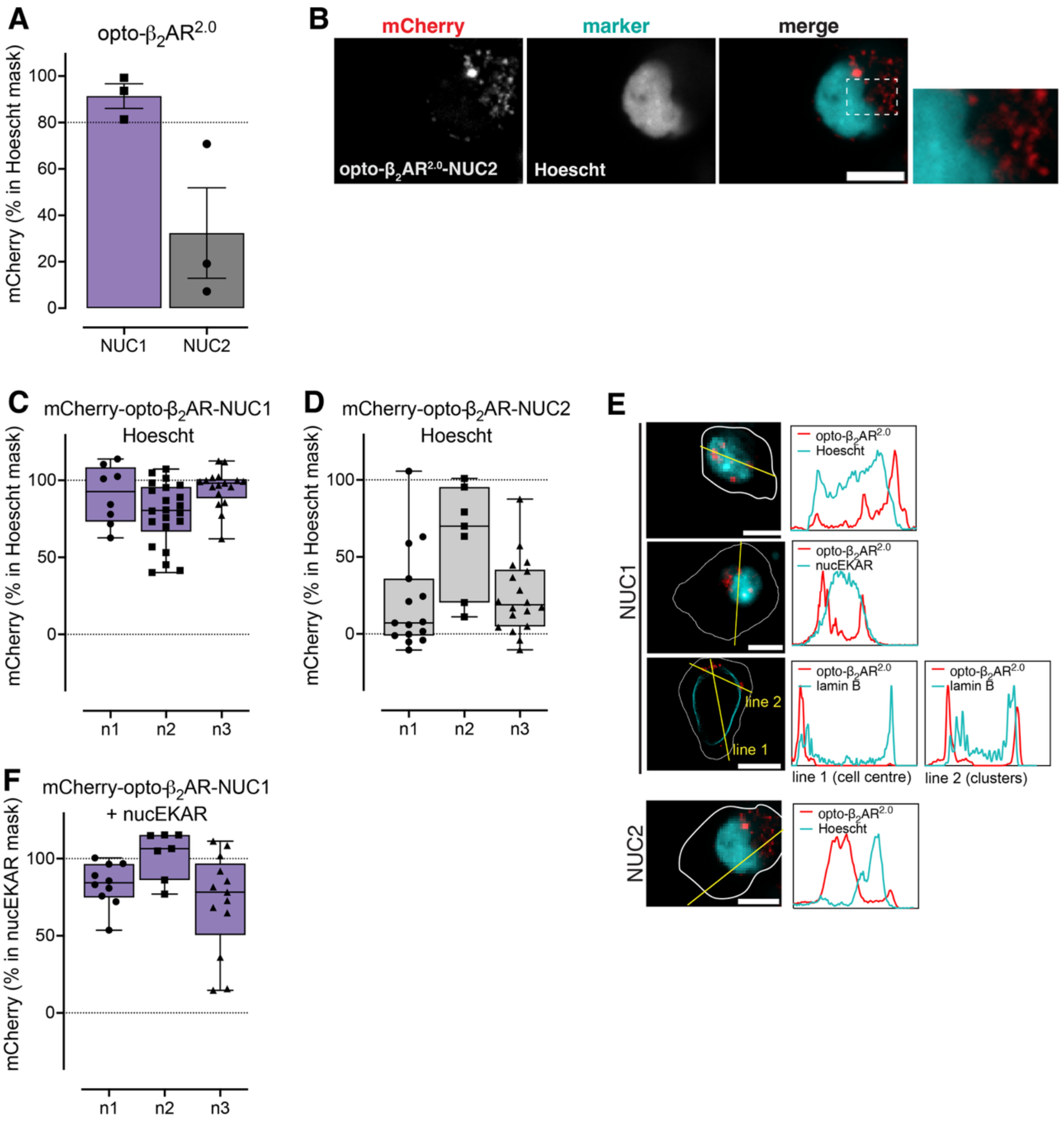


**Supplementary Figure 5. C-terminal sequence NUC1, but not NUC2, targeted the opto-$\beta_2$AR$^{2.0}$ to the nucleus.** HEK293 cells were transfected with the targeted mCherry-opto-$\beta_2$AR$^{2.0}$ and either stained with Hoescht, co-transfected with a nucelar FRET-based biosensor (EKAR), or immunostained with an antibody against an organelle resident protein. **A**) An initial screen compared the amount of mCherry fluorescence within the pixel area defined by the Hoescht stain following C-terminal fusion of NUC1 or NUC2 sequences to mCherry-opto-$\beta_2$AR$^{2.0}$. Symbols show median values from all cells from each biological replicate (shown in C and D) the bar shows the mean and error bars show SEM (n=3). **B**) Representative images show the NUC2 sequence does not localise the mCherry-opto-$\beta_2$AR$^{2.0}$ (left panel) to Hoescht (middle panel). Right panel shows merged image, and the inset (far right) shows the region boxed in white. Scale bar is 10 μm. **C-D**) The amount of mCherry-opto-$\beta_2$AR$^{2.0}$ fluorescence within the pixel area defined by the Hoescht stain was calculated for each cell in the image, following C-terminal fusion of the NUC1 (C) or NUC2 (D) sequences. Symbols show data from individual cells, bars show median and error bars show the minimum/maximum values (n=3). **E**) Line scan analysis of the representative merged images of the mCherry-opto-$\beta_2$AR$^{2.0}$ and organelle markers as displayed in B (this figure) and Figure 3D. The cell outline (white line, defined by brightfield image) was used to calculate the centroid, through which the yellow line was drawn for line scan intensity analysis (inset graphs). When this line did not sufficiently capture the organelle marker, a second line (line 2) was drawn through a region of high intensity of the marker by the blinded investigator. Scale bar is 10 μm. **F**) The amount of mCherry-opto-$\beta_2$AR$^{2.0}$-NUC1 fluorescence within the pixel area defined by the nucEKAR biosensor. Symbols show data from individual cells, bars show median and error bars show the minimum/maximum values (n=3).

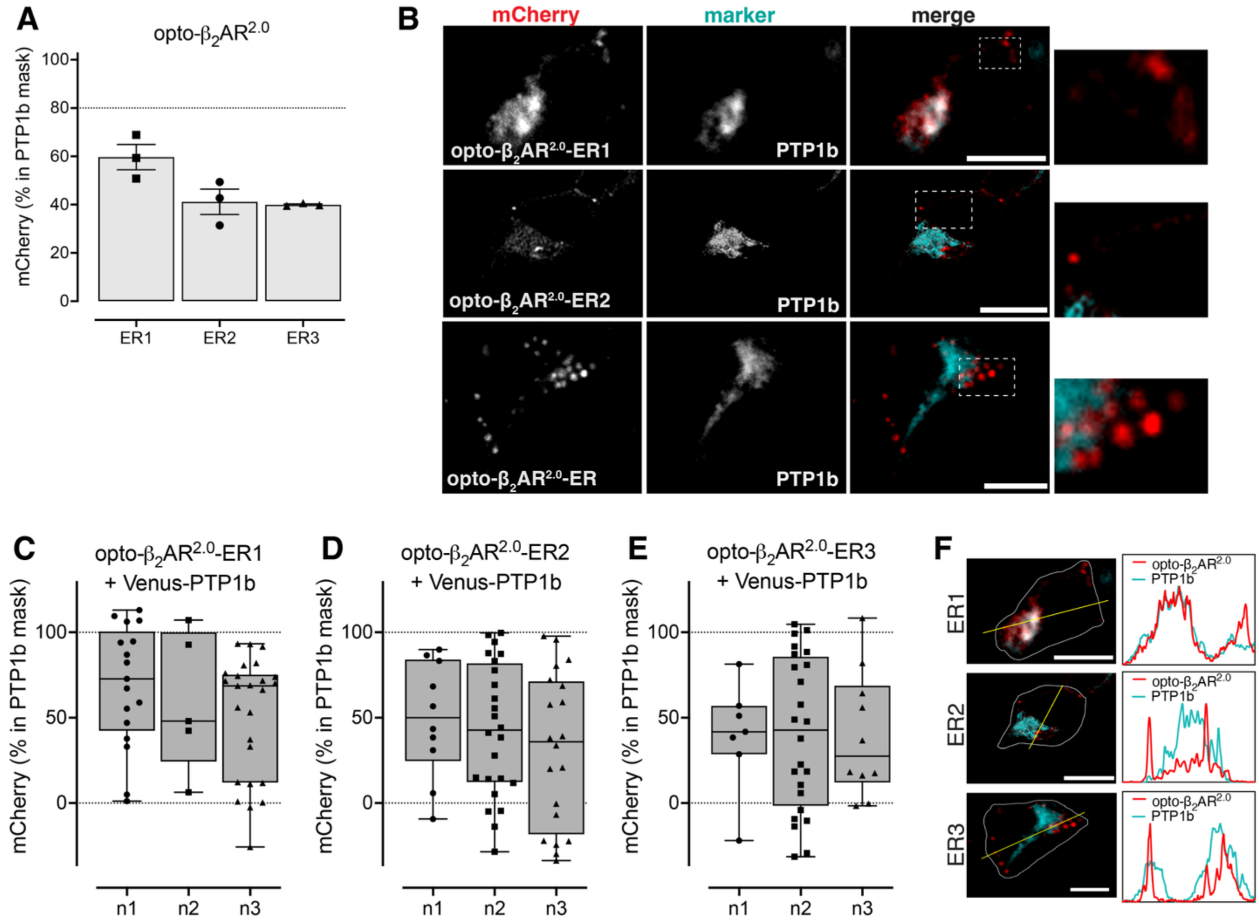


**Supplementary Figure 6. C-terminal sequences were unable to target the opto-$\beta_2AR^{2.0}$ to the endoplasmic reticulum.** HEK293 cells were co-transfected with the targeted mCherry-opto-$\beta_2AR^{2.0}$ and a Venus-PTP1b marker of the endoplasmic reticulum. **A**) An initial screen compared the amount of mCherry fluorescence within the pixel area defined by the PTP1b marker following C-terminal fusion of ER1, ER2, or ER3 sequences to mCherry-opto-$\beta_2AR^{2.0}$. Symbols show median values from all cells from each biological replicate (shown in C-E), the bar shows the mean and error bars show SEM (n=3). **B**) Representative images show none of these sequences selectively target the mCherry-opto-$\beta_2AR^{2.0}$ (left panel) to PTP1b (middle panel). Right panel shows merged image, and the inset (far right) shows the region boxed in white. Scale bar is 10 μm. **C-E**) The amount of mCherry-opto-$\beta_2AR^{2.0}$ fluorescence within the pixel area defined by the PTP1b marker was calculated for each cell in the image, following C-terminal fusion of the ER1 (C), ER2 (D), or ER3 (E) sequences. Symbols show data from individual cells, bars show median and error bars show the minimum/maximum values (n=3). **F**) Line scan analysis of the representative merged images of the mCherry-opto-$\beta_2AR^{2.0}$ and organelle marker as displayed in B. The cell outline (white line, defined by brightfield image) was used to calculate the centroid, through which the yellow line was drawn for line scan intensity analysis (inset graphs). Scale bar is 10 μm.

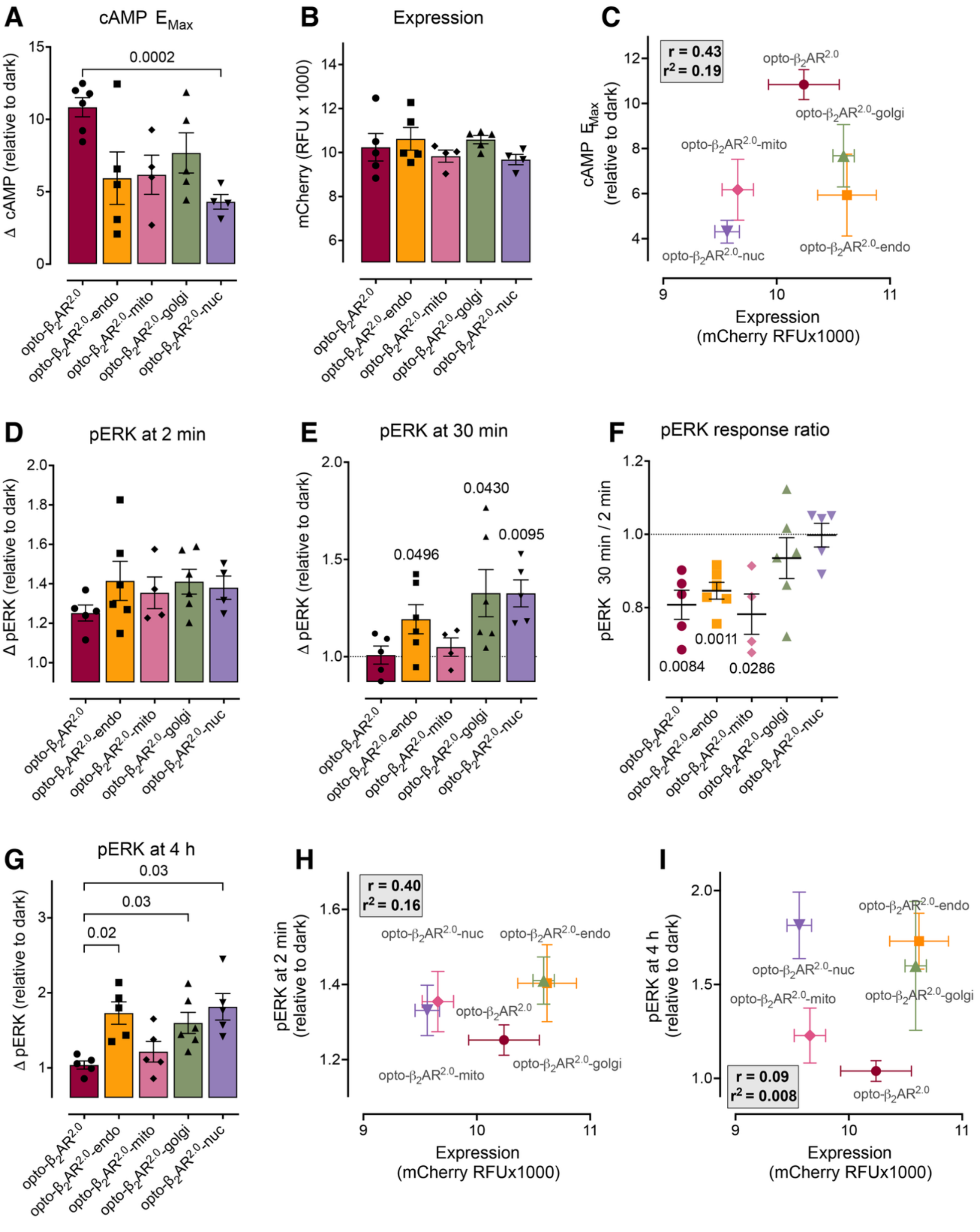


**Supplementary Figure 7. Comparison of expression level and signalling magnitude for untargeted vs targeted opto-$\beta_2$AR$^{2.0}$s.** **A**) The targeted opto-$\beta_2$AR$^{2.0}$s showed similar magnitude of cAMP responses in response to light, with opto-$\beta_2$AR$^{2.0}$-nuc inducing a smaller cAMP response than the untargeted opto-$\beta_2$AR$^{2.0}$ (n=4-6). Symbols show values from individual experiments, bars show mean and error bars show SEM; p values calculated by one-way ANOVA with Dunnett's multiple comparisons test. **B**) All opto-$\beta_2$AR$^{2.0}$s showed a similar level of mCherry fluorescence (RFU, relative fluorescence units) (n=4-5). No significant difference compared to the untargeted opto-$\beta_2$AR$^{2.0}$ as determined by one-way ANOVA with Dunnett's multiple comparisons test. Symbols show values from individual experiments, bars show mean and error bars show SEM. **C**) There was no correlation between expression level and the magnitude of the cAMP response. Symbols show means and error bars show SEM; insets show r and $r^2$ values. **D**) All opto-$\beta_2$AR$^{2.0}$s showed a similar increase in ERK phosphorylation at 2 min. No significant difference compared to the untargeted opto-$\beta_2$AR$^{2.0}$ as determined by one-way ANOVA with Dunnett's multiple comparisons test. Symbols show values from individual experiments, bars show mean and error bars show SEM. **E**) After 30 min light stimulation, only opto-$\beta_2$AR$^{2.0}$-endo, opto-$\beta_2$AR$^{2.0}$-golgi and opto-$\beta_2$AR$^{2.0}$-nuc showed elevated ERK phosphorylation compared to dark controls (n=4-6). Symbols show values from individual experiments, bars show mean and error bars show SEM; p values calculated by by one-sample t-test compared to a theoretical mean of 1. **F**) The untargeted opto-$\beta_2$AR$^{2.0}$, opto-$\beta_2$AR$^{2.0}$-endo and opto-$\beta_2$AR$^{2.0}$-mito induced transient increases in ERK phosphorylation, as indicated by a response ratio (fold change in the response at 30 min compared to the response at 2 min) that was significantly less than 1. In contrast, opto-$\beta_2$AR$^{2.0}$-golgi and opto-$\beta_2$AR$^{2.0}$-nuc showed a sustained increase in ERK phosphorylation over the 30 min time course. Symbols show response ratios for individual experiments, horizontal bar shows the mean, and error bars show SEM; p-values calculated by one-sample t-test compared to a theoretical mean of 1. **G**) After 4 h light stimulation, opto-$\beta_2$AR$^{2.0}$-endo, opto-$\beta_2$AR$^{2.0}$-golgi and opto-$\beta_2$AR$^{2.0}$-

nuc showed elevated ERK phosphorylation compared to the untargeted opto-β$_2$AR (n=4-6). Symbols show values from individual experiments, bars show mean and error bars show SEM; p values calculated by one-way ANOVA with Dunnett's multiple comparisons test. **H-I**) There was no correlation between expression level and the magnitude of ERK phosphorylation after 2 min (H) or 4 h (I) light stimulation. Symbols show means and error bars show SEM; insets show r and r$^2$ values.

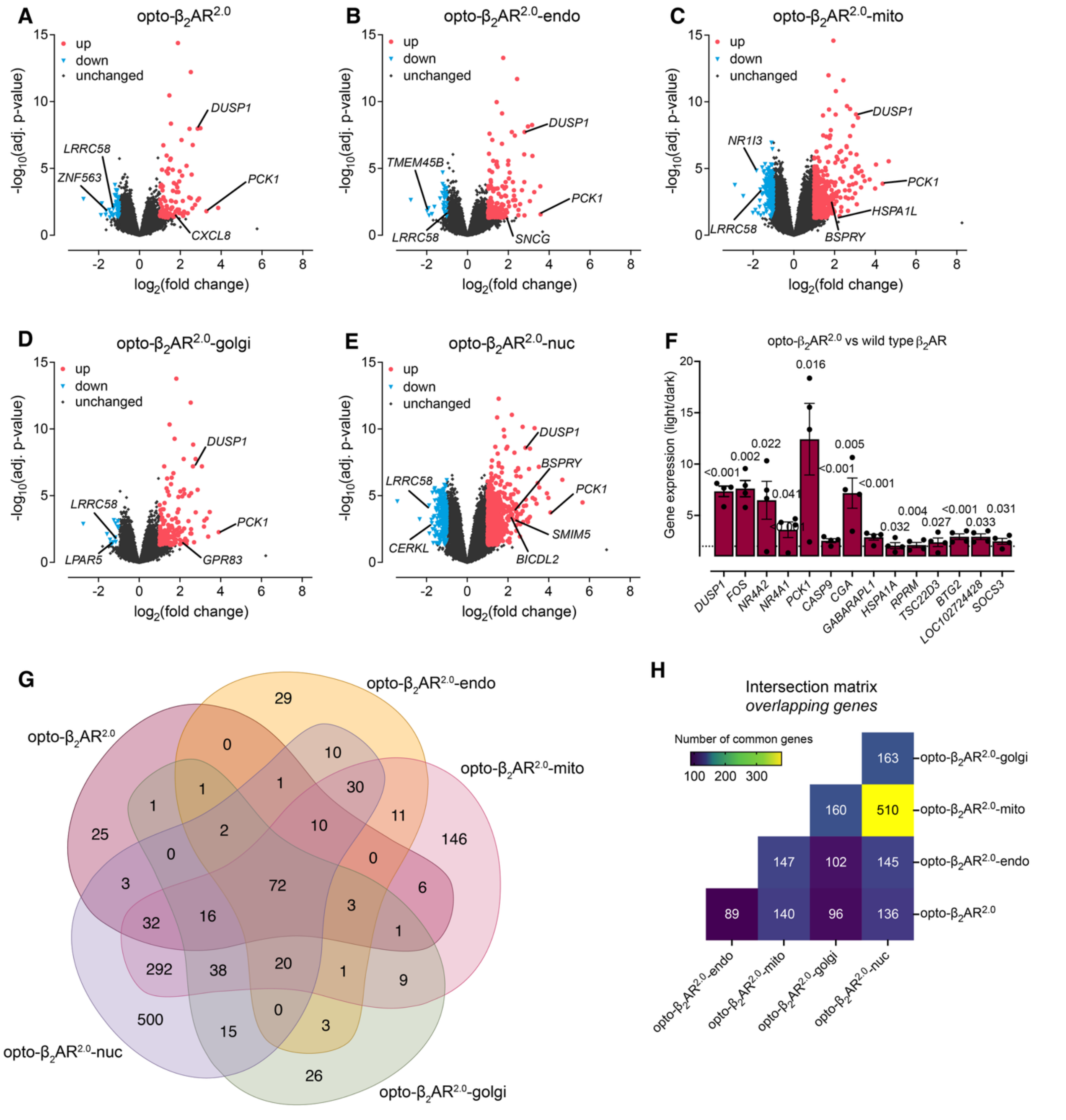


**Supplementary Figure 8. Activation of the opto-β$_2$AR$^{2.0}$ at different cellular locations leads to unique transcriptional responses.** Volcano plots showing significantly up- and down-regulated genes ($\log_2$fold-change >1, adjusted p-value <0.05) following light activation of HEK293 cells stably transfected with **A**) opto-β$_2$AR$^{2.0}$, **B**) opto-β$_2$AR$^{2.0}$-endo, **C**) opto-β$_2$AR$^{2.0}$-mito, **D**) opto-β$_2$AR$^{2.0}$-golgi, or **E**) opto-β$_2$AR$^{2.0}$-nuc (n=4). Symbols are the average over four independent experiments. Highlighted genes correspond to the top unique up- and down-regulated genes for each location (shown in Figure 6G), and the genes highlighted in Figures 6B-F if there was a significant change for that particular location. **F**) Genes significantly changed following light activation of opto-β$_2$AR$^{2.0}$ that are also regulated by ligand activation of the wild-type β$_2$AR (*10, 50*). Symbols show data expressed as the fold-change from independent experiments, bars show the mean, and error bars show SEM. Adjusted p-values are indicated above the bars. **G**) Venn diagram showing unique and commonly regulated genes across all locations. **H)** Intersection matrix showing the number of genes that were commonly regulated in response to light across two different locations.
